# Emergent RNA-RNA interactions can promote stability in a nascent phototrophic endosymbiosis

**DOI:** 10.1101/2021.04.11.439338

**Authors:** Benjamin H. Jenkins, Finlay Maguire, Guy Leonard, Joshua D. Eaton, Steve West, Benjamin E. Housden, S. Milner David, Thomas A. Richards

## Abstract

Eukaryote-eukaryote endosymbiosis was responsible for the spread of chloroplast (plastid) organelles. Stability is required for the metabolic and genetic integration that drives the establishment of new organelles, yet the mechanisms which act to stabilise nascent endosymbioses – between two fundamentally selfish biological organisms – are unclear. Theory suggests that enforcement mechanisms, which punish misbehaviour, may act to stabilise such interactions by resolving conflict. However, how such mechanisms can emerge in a nascent endosymbiosis has yet to be explored. Here, we propose that endosymbiont-host RNA-RNA interactions, arising from digestion of the endosymbiont population, can result in a cost to host growth for breakdown of the endosymbiosis. Using the model nascent endosymbiosis, *Paramecium bursaria – Chlorella* spp., we demonstrate that this mechanism is dependent on the host RNA-interference (RNAi) system. We reveal through small RNA (sRNA) sequencing that endosymbiont-derived mRNA released upon endosymbiont digestion can be processed by the host RNAi system into 23-nt sRNA. We predict multiple regions of shared sequence identity between endosymbiont and host mRNA, and demonstrate through delivery of synthetic endosymbiont sRNA that exposure to these regions can knock-down expression of complementary host genes, resulting in a cost to host growth. This process of host gene knock-down in response to endosymbiont-derived RNA processing by host RNAi factors, which we term ‘RNAi-collisions’, represents a mechanism which can promote stability in a nascent eukaryote-eukaryote endosymbiosis. By imposing a cost for breakdown of the endosymbiosis, endosymbiont-host RNA-RNA interactions may drive maintenance of the symbiosis across fluctuating ecological conditions and symbiotic status.

**SIGNIFICANCE STATEMENT:** Stable endosymbiosis between eukaryotic microbes has driven the evolution of further cellular complexity. Yet the mechanisms which can act to stabilise a nascent eukaryote-eukaryote endosymbiosis are unclear. Using the model nascent endosymbiotic system, *Paramecium bursaria–Chlorella*, we demonstrate that endosymbiont-host RNA-RNA interactions can drive a cost to host growth upon endosymbiont digestion, punishing the host for misbehaviour. These RNA-RNA interactions are facilitated by the host RNA-interference system. For endosymbiont mRNA sharing a high-level of sequence identity with host transcripts, this process can result in host gene knock-down. We propose that these endosymbiont-host RNA-RNA interactions–‘RNAi collisions’–represent a viable enforcement mechanism to sanction the host for breakdown of the endosymbiosis, promoting the stability of a nascent endosymbiotic interaction.

## MAIN TEXT

A nascent endosymbiosis between eukaryotes was a pre-requisite for the spread of photosynthetic organelles, such as plastids^1–5^. This transition from transiently engulfed cells to obligate organelles is driven by metabolic and genetic integration. Yet, to become manifest, a stable intermediary state must exist upon which this process of integration can proceed^1,3,5–8^. Conflict is an inevitable outcome of all symbioses, the resolution of which can significantly impact the stability of an interaction. Enforcement mechanisms which punish misbehaviour can act to stabilise symbioses^9,10^, yet we know little about how these can emerge in a nascent endosymbiotic interaction. *Paramecium bursaria* – a ciliate protist which harbours a clonal population of intracellular green algae, *Chlorella* spp.^11–13^ – represents a tractable model system to study emergent mechanisms in a nascent endosymbiosis^14^. The interaction is facultative^11,14–17^ and based on two-way metabolic exchange^18–25^. Endosymbiotic algae are housed within modified host phagosomes called perialgal vacuoles, which may be fused with host lysosomes to trigger digestion^26^ allowing *P. bursaria* to maintain control over the interaction in the event of conflict^27–29^. However, it is unclear how this endosymbiotic system is protected from over-exploitation by the host which would ultimately lead the interaction to collapse^30–35^.

RNA-RNA interactions can play a role in host-pathogen symbiotic systems^36–40^, whereby RNA can ‘hi-jack’ the small interfering RNA (siRNA) pathway of the symbiotic partner to modulate expression of genes involved in virulence or alternatively resistance. Whether analogous RNA-RNA interactions could occur in an endosymbiotic system has yet to be elucidated. The presence of a functional siRNA pathway in *Paramecium*, and its role in RNA-interference (RNAi), has been validated as a tool for gene silencing in *both Paramecium bursaria*^41^ and non-photo-endosymbiotic congener, *Paramecium tetraurelia*^42–46^. RNAi can be initiated through the provision of bacterial food transformed to express double-stranded RNA (dsRNA) with high sequence similarity to a target transcript^41,43^. *Paramecium* can also process single-stranded RNA (ssRNA) – including ribosomal RNA (rRNA) and messenger RNA (mRNA) – derived from a prokaryotic cell acquired through phagotrophy^42^. This processing is mediated by conserved RNAi protein components which also function in endogenous transcriptome regulation^41,42,44,47^. Significantly, these studies propose that mRNA, derived both exogenously^41,42^ and endogenously^41,44^, can act as substrates for siRNA generation in *Paramecium*.

We demonstrate that RNA released upon digestion of the algal endosymbiont is processed by the host RNAi system in *P. bursaria*. For endosymbiont-derived mRNA sharing a high-level of sequence identity with host transcripts, this processing may interfere with endogenous host gene expression resulting in a cost to host growth **(Fig. S1).** We track the interaction through sRNA sequencing, recapitulate the effect through exposure to synthetic endosymbiont RNA, and demonstrate that this mechanism is mediated by host Dicer, Piwi, Pds1 and RdRP proteins. This process of host gene knock-down in response to endosymbiont-derived RNA processing by host RNAi factors, which we term ‘RNAi-collisions’, represents a candidate enforcement mechanism which can promote stability in a nascent eukaryote-eukaryote endosymbiosis. By imposing a cost for breakdown of the endosymbiosis, endosymbiont-host RNA-RNA interactions may drive maintenance of the symbiosis across fluctuating ecological conditions and symbiotic status.

### Endosymbiont digestion in P. bursaria results in an RNAi-mediated ‘physiological cost’ to the host

*P. bursaria* can be purged of endosymbiotic algae via treatment with the ribosomal translational inhibitor, cycloheximide^14^. A comparison of ribosomal protein (RP) L29A predicted protein sequences, the active site of cycloheximide function, confirmed that *Paramecium* possess a specific nucleotide polymorphism identified as a determinant of cycloheximide resistance in other species **(Fig. S2).** This substitution was absent in all green algal species assessed, including known algal endosymbionts of *P. bursaria*. Upon treatment with cycloheximide, a significant reduction in algal-chlorophyll fluorescent intensity per host cell was observed after two-to-three days **(Fig. 1A).** A clear de-coupling of host *P. bursaria* cell number and algal fluorescence was also observed, consistent with translational inhibition in the algae but not the host **(Fig. S2),** demonstrating that loss of a photosynthetic endosymbiont population does not immediately result in a decline in host cell number. Additional staining with LysoTracker Green to identify acidic vacuoles, including lysosomes, revealed that this loss of algal fluorescence was linked to increased lysosomal activity in the host cytoplasmic environment **(Fig. 1B).** These data support that elimination of the endosymbiotic algal population during cycloheximide treatment^14,26^ is triggered by host digestion.

**Figure 1.**
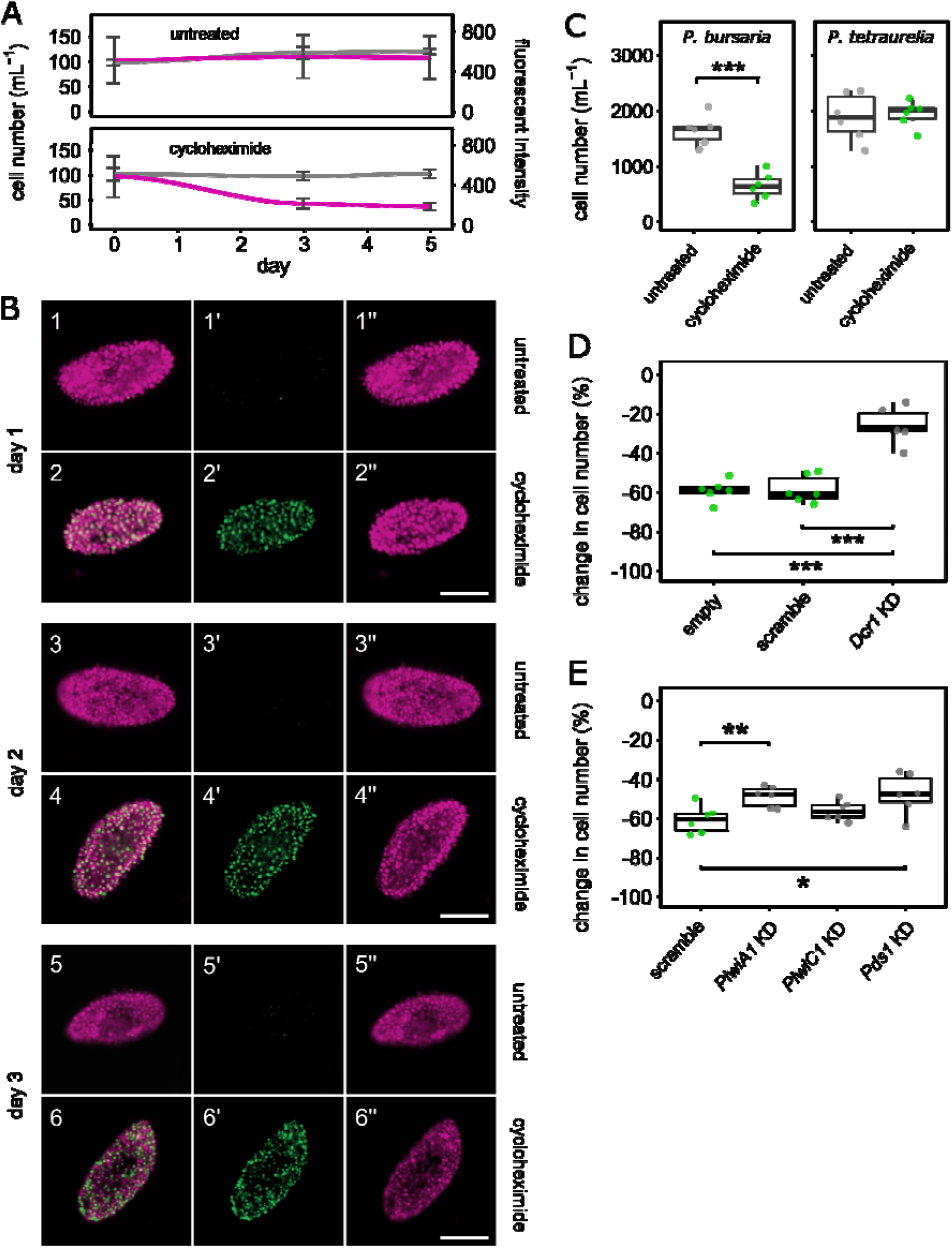
Endosymbiont digestion in *P. bursaria* results in an RNAi-mediated ‘physiological cost’ to the host. **(A)** *Paramecium* cell number (grey) vs algal chlorophyll fluorescent intensity (pink) in stationary phase *P. bursaria* cultures treated with cycloheximide (50 μgmL^-1^), compared to untreated controls. Loss of algal fluorescent intensity indicates algal death in response to cycloheximide treatment. Data are represented as mean ± SD of six biological replicates. **(B)** Stages of endosymbiont elimination in *P. bursaria* cells over three days of cycloheximide treatment (50 μgmL^-1^). Algal chlorophyll fluorescence (pink) highlights endosymbiotic algae within the *P. bursaria* cell. LysoTracker Green fluorescence (green) indicates increased host lysosomal activity in response to cycloheximide treatment. Scale bar – 50 μm. **(C)** *Paramecium* cell number in *P. bursaria* or *P. tetraurelia* cultures after 8 days of treatment with cycloheximide (50 μgmL^-1^; green), compared to untreated controls (light grey). *Paramecium* cultures were fed with *E. coli* transformed with an empty RNAi vector. **(D)** Percentage change in *P. bursaria* cell number in cultures treated with cycloheximide (50 μgmL^-1^), compared to untreated controls, after 12 days of feeding with *E. coli* expressing; Dicer (*Dcr1*) dsRNA (dark grey) to induce knock-down (KD), non-hit ‘scramble’ dsRNA or an empty vector control (green). The relative effect of Dicer dsRNA exposure indicates partial rescue of *P. bursaria* culture growth retardation in response to cycloheximide induced endosymbiont digestion. **(E)** Percentage change in *P. bursaria* cell number in cultures treated with cycloheximide (50 μgmL^-1^), compared to untreated controls, after 12 days of feeding with *E. coli* expressing; *PiwiA1, PiwiC1* or *Pds1* dsRNA (dark grey) to induce knock-down (KD), or a non-hit ‘scramble’ dsRNA control (green). **(C-E)** Feeding was conducted daily for four days prior to cycloheximide treatment, and continued throughout. Boxplot data are represented as max, upper quartile (Q3), mean, lower quartile (Q1) and min values of six biological replicates. Individual data points are shown. Significance calculated as *p ≤ 0.05, **p ≤ 0.01, and ***p ≤ 0.001, using a generalized linear model with quasi-Poisson distribution. See also **Figure S3** for the raw count data used to calculate the percentage change in cell number presented in **1D-E.**

Continued treatment with cycloheximide resulted in a significant retardation to *P. bursaria* culture growth, however this same effect was not observed in the non-photo-endosymbiotic congener species, *P. tetraurelia* **(Fig. 1C).** Both *Paramecium* species harbour the same cycloheximide resistance conferring substitution in RPL29A **(Fig. S2),** suggesting that elimination of endosymbiotic algae is costly to *P. bursaria* culture growth. Loss of algal derived photosynthate and plastid derived metabolites represents an obvious cost. However, we sought to explore whether a part of this cost may be attributed to host exposure to endosymbiont-derived RNA during digestion of the endosymbiont population, and whether this cost was mediated by the host RNAi system.

Knock-down of *Dcr1* (a host-encoded endoribonuclease Dicer required for siRNA generation) through complementary dsRNA exposure significantly rescued the cost to *P. bursaria* culture growth associated with cycloheximide treatment **(Fig. 1D** and Fig. S3). This was consistent with *Dcr1* (Dicer) knock-down in a prior study attenuating the effect of *E. coli* vector-based RNAi feeding in *P. bursaria*^41^. In the aforementioned study, knock-down of host-encoded *PiwiA1, PiwiC1* (AGO-Piwi effectors required for targeted RNA cleavage) and *Pds1* (a *Paramecium*-specific component with an unknown but essential role in exogenously induced RNAi) also attenuated an *E. coli-vector* feeding-based RNAi effect^41^. Here, knock-down of host-encoded *PiwiA1* and *Pds1* similarly rescued the cost to *P. bursaria* culture growth associated with cycloheximide treatment **(Fig. 1E** and Fig. S3). Involvement of *Pds1* confirms that the observed RNAi effect is mediated by the host, as no identifiable homologue of *Pds1* could be identified in the green algal genomes and transcriptomes sampled^41,48^. Furthermore, it excludes the possibility that off-target effects arising from host Dicer knock-down, including potential compensatory function of additional Dicer or Dicerlike paralogues in *P. bursaria*^41^, could be responsible for the RNAi effect observed. The near complete rescue observed upon partial knock-down of Dicer suggests that host Dicer processing, rather than loss of algal derived metabolites, is a more significant factor relating to growth retardation under these conditions. Taken together, these data suggest that the physiological cost to *P. bursaria* growth incurred during cycloheximide treatment, in which the endosymbiotic algae are being broken down and digested by the host, is mediated by host-encoded RNAi components. These data support the occurrence of RNAi-mediated RNA-RNA interactions between endosymbiont and host.

### Endosymbiont breakdown triggers an abundance of Dicer-dependent endosymbiont derived sRNA within *P. bursaria*

To investigate the occurrence of RNA-RNA interactions between endosymbiont and host, we tracked the abundance of endosymbiont-derived, host-processed sRNA in *P. bursaria* during endosymbiont digestion. Disruption of host RNAi was achieved through partial knock-down of Dicer, allowing us to directly test for an increase in endosymbiotic algal-derived 23-nt sRNA (the size associated with host Dicer processing^41,42,49^) resulting from cycloheximide induced endosymbiont breakdown. Upon treatment with cycloheximide, we identified an increased abundance in all 21-29 nt reads mapping to endosymbiont-derived mRNA over two-to-three days **(Fig. 2A** and **Fig. S4).** This same trend was observed for reads mapping to algal endosymbiont rRNA-derived sRNA **(Fig. S5).** Reads were mapped with 100% identity, allowing no mismatches. Any reads that additionally mapped to the host with 100% identity were removed, to ensure that the subset of sRNA detected was of definitive algal origin. A significant increase in algal mRNA-derived 23-nt sense and antisense sRNA demonstrates a greater abundance of potential RNAi substrates during endosymbiont digestion^41,42,49^.

**Figure 2.**
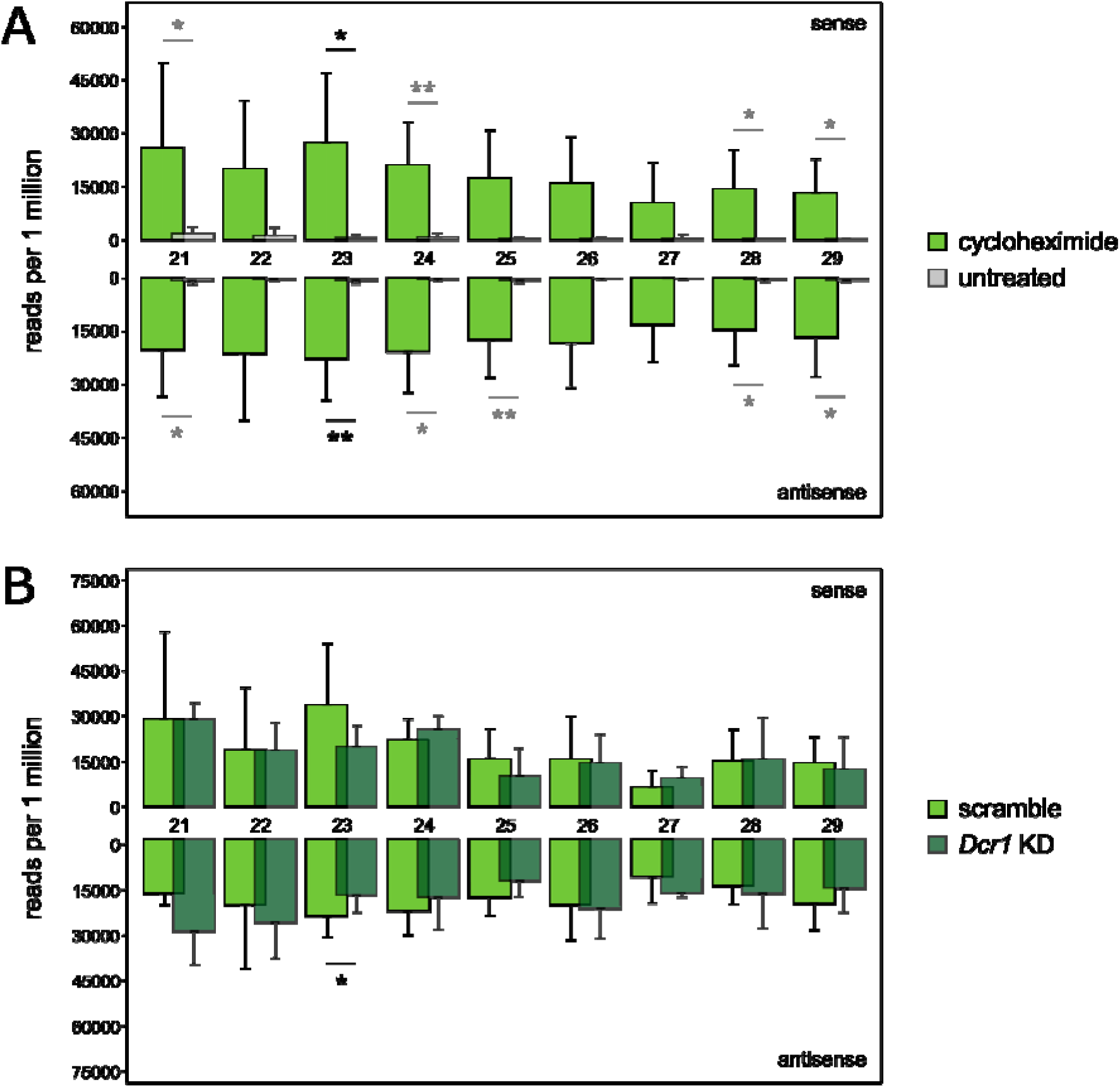
Endosymbiont breakdown triggers an influx of Dicer-dependent endosymbiont-derived sRNA within *P. bursaria*. **(A)** Size distribution (nt) of sRNA mapped to endosymbiont-derived cytoplasmic mRNA. sRNA was extracted from *P. bursaria* cultures over two days (day 2 and 3) of cycloheximide treatment (50 μgmL^-1^), or from untreated controls. Note the relative increase in 23-nt abundance between untreated and cycloheximide treated cultures. *P. bursaria* cultures were fed with *E. coli* transformed to express non-hit ‘scramble’ dsRNA. Data are represented as mean ± SD of six biological replicates, and normalised against total endosymbiont mRNA-mapping 21-29-nt reads per dataset. (B) Size distribution (nt) of sRNA mapped to endosymbiont-derived cytoplasmic mRNA. sRNA was extracted from *P. bursaria* cultures on day 3 of cycloheximide treatment (50 μgmL^-1^). *P. bursaria* cultures were fed with *E. coli* transformed to express Dicer (*Dcr1*) dsRNA to induce knock-down (KD), or a non-hit ‘scramble’ dsRNA control. Note the relative increase in 23-nt abundance during cycloheximide treatment in *P. bursaria* cultures exposed to scramble dsRNA, compared to Dicer dsRNA. Data are represented as mean ± SD of three biological replicates, and normalised against total endosymbiont mRNA-mapping 21-29-nt reads per dataset. (**A-B**) Feeding was conducted daily for four days prior to cycloheximide treatment, and continued throughout. Significance calculated as *p ≤ 0.05, **p ≤ 0.01, and ***p ≤ 0.001 using a generalised linear model with quasi-Poisson distribution. All curated ‘endosymbiont’ mRNA transcript bins used for sRNA mapping are available on Figshare (10.6084/m9.figshare. 12301736). See also **Figure S4** for sRNA abundance at each individual day.

Partial knock-down of host Dicer during cycloheximide treatment significantly ablated this endosymbiont mRNA-derived 23-nt antisense abundance after three days of cycloheximide treatment **(Fig. 2B** and **Fig. S4).** This suggests that an increase in endosymbiont mRNA-derived 23-nt antisense sRNA during endosymbiont digestion is dependent on host Dicer function. This is consistent with knock-down of host Dicer specifically reducing 23-nt sRNA abundance in Paramecium^41,42,49^, and the observation that *Paramecium* RNAi factors are capable of processing both endogenously and exogenously derived mRNA^42,44^. It is important to note that this effect likely under-represents the full extent of host Dicer processing, due to the requirement for a paradoxical and therefore incomplete Dicer perturbation through Dicer-dependent RNAi-based knock-down^41^. Furthermore, these data represent only a subset of the sRNA potentially present due to the stringent mapping approach used to identify endosymbiont-derived sRNA, as any reads that additionally mapped to host RNA template with 100% sequence identity in either orientation were excluded from this analysis. While these reads with high shared sequence identity are the most important subset of sRNA for the identification of putative mRNA-mRNA interactions between endosymbiont and host, their exclusion here has allowed us to identify host RNAi processing of definitively endosymbiont-derived transcripts. Interestingly, one of the host-derived transcripts that mapped with 100% identity to an excluded endosymbiont-derived 23-nt sRNA was *P. bursaria* heat shock protein 90 (*HSP90*). An investigation of regions of high shared sequence identity between endosymbiont and host is explored below.

Next, we conducted a series of control observations. Firstly, we assessed whether cycloheximide treatment was altering general host-derived sRNA production, despite the inferred host resistance discussed above **(Fig. S2).** Reads were mapped to a dataset of 20 host transcripts that contained no potential 23-nt overlap with any identified algal transcripts (allowing for ≤2-nt mismatches), to ensure that these host transcripts were unaffected by an increased rate of putative RNA-RNA interactions derived from the endosymbiont **(Fig. S6).** No significant increase in 23-nt abundance was observed for reads mapping to this subset of host mRNA transcripts during cycloheximide treatment, consistent with host resistance to cycloheximide not altering the host-derived population of sRNAs with low sequence identity to algal mRNA. Secondly, we assessed whether endosymbiotic algal strains cultured under free-living conditions would also generate an abundance of 23-nt sRNA upon cycloheximide treatment **(Fig. S7).** Importantly, no clear increase in 21-25 nt algal-derived sRNA abundance was observed during algal treatment with cycloheximide when grown outside of the host cytoplasmic environment, nor was a 23-nt peak in algal derived sRNAs evident. This is consistent with data from a previous study which found that the algal endosymbiont of *P. bursaria* was not actively generating sRNA >20-nt^41^. These results suggest that increased algal endosymbiont-derived 23-nt sRNA abundance upon treatment with cycloheximide is dependent on host sRNA processing within the endosymbiotic system. Taken together, these data indicate that an abundance of endosymbiont-derived RNAs are released during endosymbiont digestion, which then act as substrate for processing by the host RNAi system, resulting in 23-nt sRNAs. Importantly, processing of endosymbiont-derived sRNA in a host Dicer-dependent manner indicates that these interactions are occurring in the host cytoplasm, supporting the hypothesis that endosymbiont-host mRNA-mRNA interactions are possible.

### Comparison of transcriptome data reveals the potential for host transcript interaction by endosymbiont-derived RNAs

To understand the extent of possible RNA-RNA interactions between endosymbiont and host, we built a bioinformatic transcriptome processing tool, eDicer (https://github.com/fmaguire/eDicer), which allows identification of all possible Dicer-generated sense and antisense oligonucleotides produced from a given transcriptome dataset. By mapping 23-nt reads to a second (‘host’) dataset allowing for ≤2-nt mismatches, potential RNA-RNA interactions between an input (endosymbiont, vector, or food) and a host RNA population can be identified. Inclusion of reads with ≤2-nt mismatches were based on the tolerance for mismatching complementarity reported during RNAi-mediated knockdown of gene expression in multiple systems^50–54^. Confirmation of a similar mismatch tolerance in *P. bursaria* is shown in **Fig. 3E & Fig. S12.** For a full overview of the eDicer comparative analysis process, please refer to the **Supplementary Methods.**

**Figure 3.**
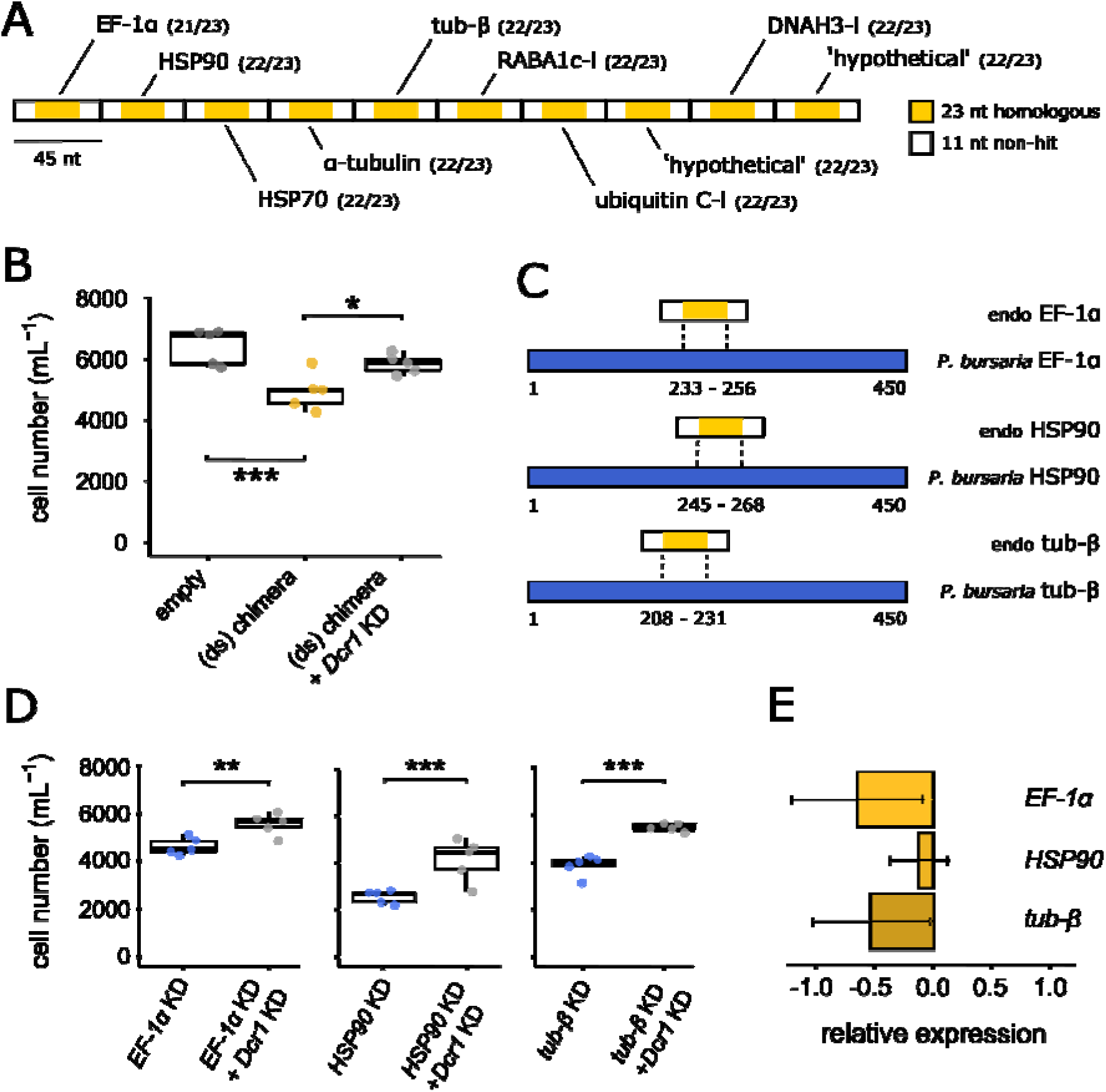
*In vivo* exposure to a synthetic endosymbiont-derived RNA chimera generates simultaneous knock-down of homologous host genes in *P. bursaria*. (**A**) Schematic showing chimeric construct design. Ten endosymbiont-derived 45-nt transcript sequences **(Table S3),** featuring a 23-nt region with >91% sequence identity between endosymbiont and host (yellow) flanked by 11-nt of ‘non-hit’ algal transcript (white). Numbers in brackets denote sequence identity between endosymbiont and host. (**B**) *P. bursaria* cell number after 12 days of feeding with *E. coli* expressing: chimera dsRNA (yellow); chimera dsRNA mixed with Dicer (*Dcr1*) dsRNA (light grey; rescue); or an empty vector control (dark grey). See **Figure S9** for an experimental dilution of dsRNA chimera delivery. (**C**) Schematic demonstrating the degree of overlap between three endosymbiont (endo) derived transcripts from the chimeric construct (white/yellow) and respective 450-nt homologous region of the host (*P. bursaria*) transcript (blue). Each region of 23-nt overlap (yellow) represents putative RNAi ‘collisions’ between endosymbiont and host. (**D**) *P. bursaria* cell number after 12 days of feeding with *E. coli* expressing: host *EF1-α, HSP90* or *tub-β* 450-nt dsRNA (blue) to induce host knock-down (KD), compared to Dicer (*Dcr1*) dsRNA mixed controls (grey; rescue phenotype). See also **Figure S11** for individual knock-down (KD) of each of the ten broader host targets of the endosymbiont-derived dsRNA chimera. (**E**) qPCR of mRNA extracted from day 3 of chimera-RNAi feeding (**3B**), revealing knock-down of *EF1-α, HSP90* and *tub-β* host gene expression in *P. bursaria* in response to endosymbiont derived chimera dsRNA exposure. Standardised expression of an Actin housekeeping gene was used for normalisation. Data are represented as mean ± SD of three biological replicates. These three genes were assessed for knock-down as a result of endosymbiont derived chimera exposure, based on evidence that directed knock-down of the wider corresponding host gene led to retardation of *P. bursaria* culture growth (**3D**). For an extended figure showing the effect of Dicer knock-down on the expression of these three host genes, see **Figure S12. (B/D)** Multiple vector delivery was conducted at a 50:50 ratio during feeding. Boxplot data are represented as max, upper quartile (Q3), mean, lower quartile (Q1) and min values of five biological replicates. Individual data points are shown. Significance for boxplot data calculated as *p ≤ 0.05, **p ≤ 0.01, ***p ≤ 0.001, using a generalized linear model with quasi-Poisson distribution.

Using a dataset consisting of transcripts binned as either ‘endosymbiont’ or ‘host’ (using a curated *P. bursaria* transcriptome^55^), we identified 35,703 distinct 23-nt putative mRNA-mRNA interactions between the *P. bursaria* ‘host’ and ‘endosymbiont’ RNA populations, representing 0.121% of the total inventory of distinct host 23-nt k-mers identified **(Fig. S8A;** see also **Table S2 & Supplementary Methods).** This was found to be 120-fold greater than the number of putative 23-nt mRNA-mRNA interactions predicted between bacterial food and host transcripts (from two different bacterial sources). Furthermore, the ratio of total: ‘potentially-lethal’ putative 23-nt mRNA-mRNA interactions (determined by cross-referencing genes known to be conditionally essential in *Saccharomyces cerevisiae*^56^) was found to be greater between endosymbiont-and-host transcripts (1:1.4) than between bacterial food-and-host transcripts (1:0.16/0.17). Similar patterns were observed for predicted rRNA-rRNA interactions **(Fig. S8B,** see also **Table S2** & **Supplementary Methods**). These *in silico* analyses demonstrate that there is far greater potential for the occurrence of both mRNA-mRNA and rRNA-rRNA interactions between the algal endosymbiont and host (eukaryote-eukaryote) RNA populations, than there is between bacterial food and host (prokaryote-eukaryote) RNA populations.

### *In vivo* exposure to a synthetic endosymbiont mRNA-derived chimera generates knock-down of high-identity host transcripts

Having identified the occurrence of putative endosymbiont-host RNA-RNA interactions *in silico, we* assessed whether exposure to synthetic fragments of algal mRNA could recapitulate the cost to host growth observed during endosymbiont digestion **(Figure 1E).** We identified ten endosymbiont mRNA interaction fragments that shared >91% (or 21-nt) sequence identity with host transcripts across a 23-nt region, selected at random from the analysis reported in **Fig. S8A.** Each mRNA interaction fragment was chosen to contain at least 1 SNP specific to the endosymbiont to ensure that any identifiable effect could be attributed to an endosymbiont-like transcript, rather than the host version. These interaction fragments were predicted to still be effective RNAi templates for *P. bursaria* based on the tolerance for mis-matching complementarity observed in RNAi-mediated knock-down in other systems^50–54^ (and confirmed below in *P. bursaria*; **Fig. 3E & Fig. S12).** Six mRNA interaction fragments showed putative homology to ‘non-lethal’ yeast genes (including *EF1-α* and *HSP90*); two showed putative homology to ‘lethal’ yeast genes (including *tub-β*); and the remaining two had no identifiable homologues in yeast **(Table S1).** Significantly, *HSP90* was the host transcript previously identified as a candidate for putative endosymbiont-host RNA-RNA interaction in the sRNA analysis of **Fig. 2.** However, to ensure that any identifiable effect could be attributed to the endosymbiont-derived transcript, the newly identified interaction fragment was chosen from a different region of this gene containing 2 SNPs specific to the endosymbiont. All ten mRNA interaction fragments were composed of the predicted interacting 23-nt sRNA sequence **(Fig. S8A)** flanked by 11-nt of contiguous ‘non-hit’ endosymbiont transcript as a filler on each side **(Table S3).** These ten 45-nt fragments were combined into a single 450-nt synthetic endosymbiotic algal chimera **(Fig. 3A),** cloned into an L4440 plasmid, and transformed into *E. coli* for feeding-induced RNAi.

Exposure to endosymbiotic algal chimera dsRNA resulted in significant retardation of *P. bursaria* culture growth **(Fig. 3B).** Once more, this effect was attenuated by knock-down of host Dicer, demonstrating an RNAi-mediated response to synthetic endosymbiont-host mRNA-mRNA interactions that resembled the host response to endosymbiont digestion in **Fig. 1C-E.** It is important to note that synthetic exposure to endosymbiont-derived RNA via an *E. coli* feeding vector would likely over-represent these putative RNA interactions. However, to address this issue, we designed a non-hit ‘nonsense’ control composed of a tandem assembly of the 11-nt regions of contiguous ‘non-hit’ algal transcript present in the chimera. A relative dilution of endosymbiont chimera dsRNA delivery alongside the ‘nonsense’ control (1, 1:1 and 1:3) also resulted in *P. bursaria* culture growth retardation **(Fig. S9),** demonstrating that a reduced relative delivery of synthetic endosymbiont RNA can also result in a cost to host growth. Similar experiments were conducted to investigate the possibility of endosymbiont-host rRNA-rRNA interactions, but showed that there was no equivalent effect on host growth, suggesting that ribosomal RNA is shielded from such effects **(Fig. S10,** see also **Table S4).**

To identify which putative mRNA-mRNA interactions were capable of facilitating a cost to *P. bursaria* culture growth, we identified the ten 450-nt host transcripts predicted to be hit by each endosymbiont derived 23-nt mRNA interaction fragment used in the chimera (**Figure 3C** - three examples are shown). Individual exposure to dsRNA corresponding to each individual but longer *P. bursaria* transcript-form identified three which resulted in a significant cost to host growth, relative to Dicer knock-down controls (**Fig. 3D;** see **Fig. S11** for all 10). These *P. bursaria* transcripts correspond to elongation factor-1α (*EF-1α*), heat shock protein 90 (*HSP90*; the host transcript identified as a candidate for putative endosymbiont-host RNA-RNA interaction in the sRNA analysis of **Fig. 2**), and tubulin-β chain (*tub-β*). It can therefore be inferred that these are the 23-nt mRNA interaction fragments **(Fig. 3A/C)** that likely resulted in a cost to host growth observed during synthetic endosymbiont chimera dsRNA exposure **(Fig. 3B)**. Using mRNA extracted from *P. bursaria* during chimera-RNAi feeding described above **(Fig. 3B),** qPCR revealed a reduction in host transcript expression of *EF-1α* and *tub-β* in response to endosymbiont chimera dsRNA exposure **(Fig. 3E)**. Host transcript expression of *HSP90* appears inconclusive, however, expression of all three host transcripts was partially rescued upon knock-down of host Dicer **(Fig. S12).** From these data we can therefore conclude that 23-nt of >91% complementary ‘endosymbiont’ transcript is sufficient to facilitate detectable knock-down of a corresponding host transcript, demonstrating that exposure to endosymbiont-derived RNA originating from phagosomes is capable of impacting host gene expression via host RNAi knock-down.

It is important to consider that delivery of a synthetic endosymbiont-derived RNA chimera represents only an approximation of the putative 23-nt RNA-RNA interactions occurring in *P. bursaria*. Exposure to endosymbiont RNA via an *E. coli* feeding vector would likely over-represent these putative interactions. However, the random generation of 23-nt Dicer substrates across the dsRNA chimera (only three regions of which resulted in a detectable cost to host growth; **Fig. 3A/D & Fig. S11)** represents only part of the wider *E. coli*-derived RNA population processed by the host RNAi system during chimera-RNAi feeding. Together with dilution of endosymbiont chimera dsRNA delivery **(Fig. S9),** these observations support the hypothesis that only a relatively small number of 23-nt RNA-RNA interactions can induce a cost to host growth in *P. bursaria*. Furthermore, these data confirm that sRNAs with ≤2-nt mismatches are effective templates for RNAi-mediated knock-down of gene expression in *P. bursaria*, suggesting that the sRNA analysis in **Fig. 2** may under-represent the true scale of endosymbiont-derived sRNAs that can result in RNA-RNA interactions between endosymbiont and host. Importantly, we note that the occurrence of putative endosymbiont-host RNA-RNA interactions does not exclude a wider cost to host growth arising from processing a large population of algal derived RNAs with no defined host target, a process which must pose a cost to host cellular economics and transcriptional control/fidelity. For further justification of this ‘synthetic’ approach, and a consideration of the results that we can reliably draw from these data, see **Discussion S1.**

### Single-stranded delivery of synthetic endosymbiont-derived RNA, analogous to endosymbiont mRNA, results in a cost to *P. bursaria* growth

As RNA derived naturally from the endosymbiont was unlikely to be double-stranded, two further constructs were designed to assess the efficacy of synthetic endosymbiont chimera single-stranded RNA (ssRNA) exposure in *P. bursaria* **(Fig. 4 & Fig. S13)**. Exposure to endosymbiont chimera ssRNA also resulted in a significant cost to *P. bursaria* culture growth, however this effect was only observed when delivered in the sense orientation ([+]ssRNA). Notably, chimera [+]ssRNA exposure resulted in a greater cost to host growth than chimera dsRNA exposure. A similar orientation bias was observed upon Dicer knock-down during cycloheximide treatment in a prior experiment, in which delivery of Dicer [+]ssRNA rescued culture growth to a greater extent than delivery of Dicer [-]ssRNA or Dicer dsRNA **(Fig. S14).** Importantly, the orientation of [+]ssRNA represents the same orientation as the mRNA transcripts from which each of these target templates were identified.

**Figure 4.**
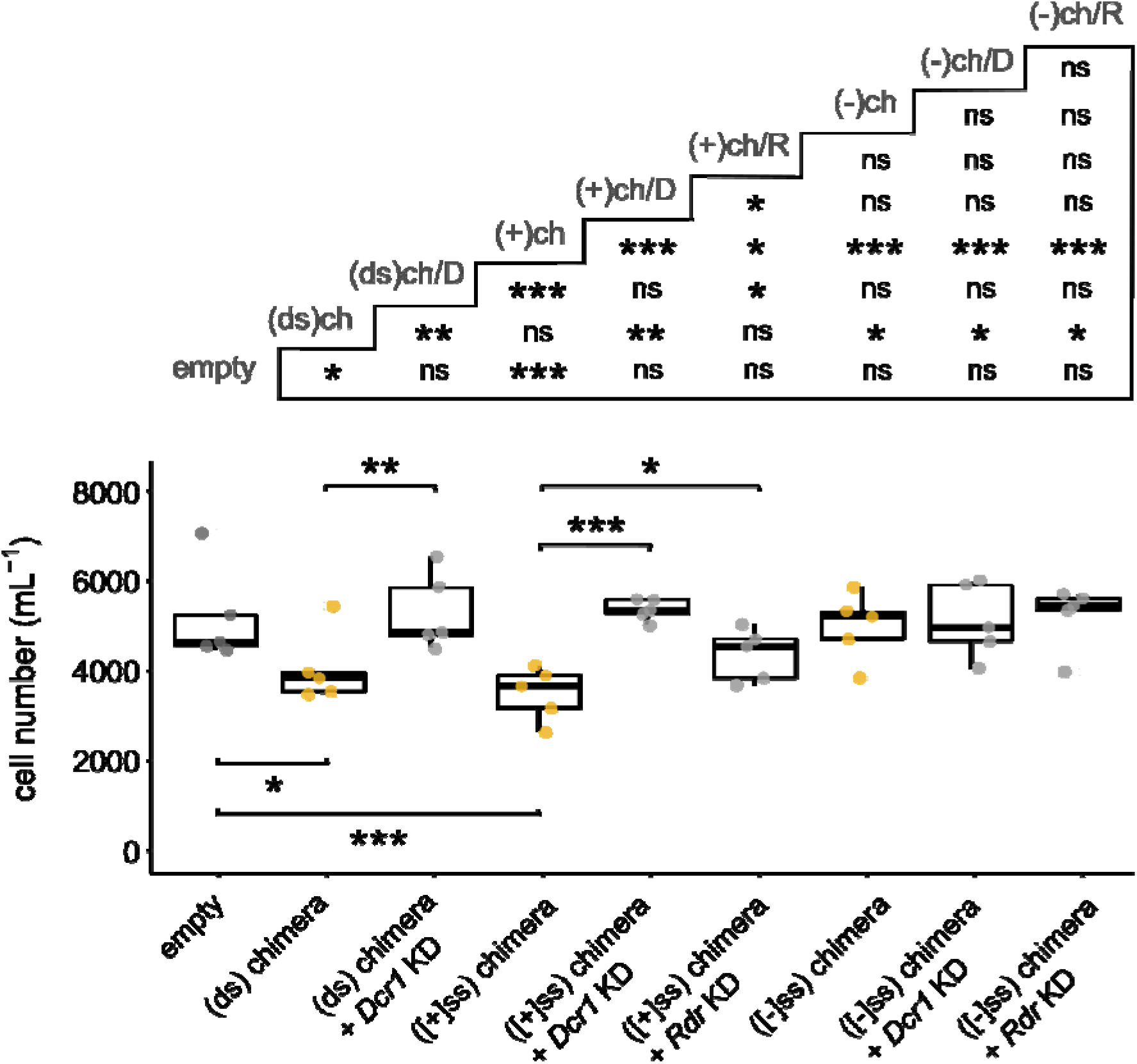
Single-stranded delivery of synthetic endosymbiont-derived RNA, analogous to endosymbiont mRNA, results in a cost to *P. bursaria* growth. *P. bursaria* cell number after 14 days of feeding with *E. coli* expressing; chimera RNA in ds, [+]ss or [-]ss orientation (yellow); chimera RNA in ds, [+]ss or [-]ss orientation mixed with Dicer (*Dcr1*) or RdRP (*Rdr*) dsRNA (light grey; rescue); or an empty vector control (dark grey). Multiple vector delivery was conducted at a 50:50 ratio during feeding. Asterisks displayed in the grid above denote pairwise significance values. Boxplot data are represented as max, upper quartile (Q3), mean, lower quartile (Q1) and min values of five biological replicates. All significance for boxplot data calculated as *p ≤ 0.05, **p ≤ 0.01, ***p ≤ 0.001, ‘ns’ no significance, using a generalized linear model with quasi-Poisson distribution.

Knock-down of host Dicer (dsRNA delivery) was able to attenuate the cost to host growth associated with endosymbiont chimera [+]ssRNA exposure **(Fig. 4).** Importantly, simultaneous knock-down of host *Rdr1* and *Rdr2* (RNA-dependent RNA polymerases involved in amplification of primary or secondary sRNA triggers during RNAi^42,46^) also significantly rescued this cost to host growth. This is consistent with the role of RdRP proteins in processing exogenous sRNA, partially degraded mRNA cleavage products, or fulllength mRNA transcripts in *Paramecium*^42,44–46^. As for Dicer, knock-down of host RdRP through RNAi represents a potential paradox, and so constitutes only partial or transient loss of function^41^. However, we are again confident that disruption in this manner would be sufficient to perturb RdRP-mediated RNAi in *P. bursaria* after 14 days, as seen in **Fig. 4.** While the mechanism of down-stream Dicer processing of RdRP-generated dsRNA substrates remains unknown (see **Discussion S1)**, we can nonetheless infer from these data that single-stranded RNA-induced RNAi knock-down in *P. bursaria* is partially dependent upon host RdRP function.

Irrespective of the mechanistic basis for a sense oriented ssRNA knock-down bias, it is important to note that delivery of chimera [+]ssRNA demonstrated here represents the same orientation as the endosymbiont derived mRNA interaction fragments from which these putative endosymbiont-host RNA-RNA interactions were identified. This is consistent with the observation that *Paramecium* RNAi factors are capable of processing both endogenously and exogenously derived mRNA^42,44^. The ability to facilitate a cost to host growth through delivery of single-stranded RNA, in the same orientation as the host target mRNA, therefore supports the hypothesis that a cost to host growth upon endosymbiont digestion can be facilitated by host RNAi-mediated RNA-RNA interactions between endosymbiont and host mRNA populations.

## DISCUSSION

Through manipulation of the *P. bursaria – Chlorella* spp. endosymbiotic system, we have demonstrated that RNA released upon digestion of the algal endosymbiont is processed by the host RNAi system. For endosymbiont-derived mRNA sharing a high-level of sequence identity with host transcripts, this processing can result in knock-down of endogenous host gene expression, resulting in a cost to host growth **(Fig. S1).** We therefore postulate that these RNA-RNA interactions are of importance in eukaryote-eukaryote endosymbioses where partners are likely to share greater sequence similarity, especially among conserved transcripts which tend to be more highly represented among lethal or conditionally essential genes^56^. Due to the inherent difficulty in characterising such mechanisms directly, we have relied on a multiple experimental approach to demonstrate the viability of putative RNA-RNA interactions at each stage of the process. We have tracked the interaction through sRNA sequencing, recapitulated the effect through exposure to synthetic endosymbiont-derived RNA – including sense ssRNA analogous to endosymbiont mRNA – and demonstrated that this mechanism is mediated by host Dicer, Piwi, Pds1 and RdRP proteins. This process of host gene knock-down in response to endosymbiont-derived RNA processing by host RNAi factors, which we term ‘RNAi-collisions’, represents a candidate enforcement mechanism to sanction the host for breakdown of the interaction, a factor that would promote stability in a nascent eukaryote-eukaryote endosymbiosis.

The long-term maintenance of symbiotic interactions represents a quandary for evolutionary theory^30–35^. How do such relationships avoid over-exploitation by one partner that would ultimately lead the interaction to collapse? Partner switching is one option, however the result is typically a reduced pattern of co-evolution between symbiont and host that can inhibit the process of metabolic and genetic integration, stalling the evolution of stable interactions which are needed if a system is to move towards evolution of an organelle^13,57–60^. Previous studies have suggested that when selfish behaviours arise, the evolution of enforcement to punish or suppress the exploitative partner can act to restore cooperation^9,10^. Enforcement mechanisms have been identified in diverse biological systems^61,62^, and are argued to be one of the most effective drivers of cooperation in egalitarian alliances between different species, such as that between *P. bursaria* and its algal endosymbiont^9^. The results presented here suggest that ‘RNAi collisions’ between endosymbiont and host, which are capable of imposing a cost to host growth for breakdown of the symbiosis, could provide an emergent enforcement mechanism to discourage over-exploitation of the endosymbiont population by the host.

Interestingly, the emergence of this mechanism appears to be a by-product of pre-existing biological features that are already likely to be under strong selective pressure. For instance, the widely functional RNAi system of the host^42,43,45^, or the conserved gene repertoire and sequence composition of the host and endosymbiont transcriptomes from which some ‘RNAi collisions’ have here been identified (including many transcripts which are conditionally essential in other systems^56^). Unlike comparable mechanisms of RNA-RNA interactions that have been studied in host-pathogen symbioses^36–40^, these ‘RNAi collisions’ appear to be untargeted and, hence, emergent. In the aforementioned host-pathogen systems, targeted RNA is passed from one partner to the other in order to modulate expression of transcripts involved in virulence or resistance^36,39,40^. However, in order for such systems to evolve, they must first exist in an untargeted form upon which selection is able to act, allowing the emergence of specific RNA factors^37,38^. Identification of undirected ‘RNAi collisions’ in the *P. bursaria – Chlorella* system represents one such intermediary state, emergent in nature and untargeted, upon which sustained cellular interaction coupled with the potential for host-symbiont conflict could drive the selection of targeted RNA-RNA interactions.

We therefore propose that ‘RNAi collisions’ represent a putative mechanism to discourage over-exploitation of the endosymbiont population by the host. Here we have used the example of mass endosymbiont digestion in response to drug treatment to simulate this effect in the extreme. In natural interactions between *P. bursaria* and its algal endosymbiont, such a cost would only need to occur in the drastic occurrence of mass endosymbiont digestion in order to drive stability of the interaction. Importantly, the endosymbiotic algal population within *P. bursaria* is largely composed of closely related or clonal lineages^11–13^, and as such, the fate of the algal population should be considered as a collective unit. This cost need only act to supress large-scale, rapid destruction by the host in order to drive the maintenance of a surviving subsection of the endosymbiont population. Previous studies have demonstrated how *P. bursaria* is capable of manipulating endosymbiont load in response to varying light conditions to better suit its own ends^27,28^, however, in these examples, reduction of endosymbiont number through digestion is slow and partial. By providing a system that selects against rapid and near-complete digestion of the endosymbiont population, ‘RNAi collisions’ effectively buffer the nascent endosymbiotic interaction against total breakdown. We suggest that this has allowed the relationship to be maintained across time and varying ecological conditions, even in the event of host-symbiont conflict^27,28^ and fluctuating endosymbiont numbers^14,26,27^. As an alternate route to conflict resolution that avoids partner switching, we propose that such a mechanism would facilitate greater co-evolution between endosymbiont and host. Over time, this would allow the metabolic and genetic integration that drives the formation of obligate symbioses to become manifest. We therefore present ‘RNAi collisions’ as a new mechanism in this endosymbiotic system – a factor which can promote stability in the face of conflict in an emergent endosymbiotic eukaryote-eukaryote cell-cell interaction.

## AUTHOR CONTRIBUTIONS

B.H.J., D.S.M., and T.A.R. conceived and designed the experiments. F.M. and T.A.R. conceived and designed the eDicer computational analysis. B.H.J., D.S.M., and T.A.R. wrote the manuscript. B.H.J., D.S.M., F.M. and G.L. conducted experimental work and analysed the data. B.E.H., S.W. and J.D.E aided in conceptual and experimental design, and in conducting experimental work.

## ACKNOWLEDGEMENTS

This work was primarily supported by an EMBO YIP award and a Royal Society University Research Fellowship (UF130382) and latterly by an ERC Consolidator Grant (CELL-in-CELL) to T.A.R. S.W. and J.D.E. were supported by awards from the Wellcome Trust (WT107791/Z/15/Z) and the Lister institute. F.M. was supported by a Donald Hill Family Fellowship in Computer Science. We thank Karen Moore and the University of Exeter Sequence Service for support with the various sequencing projects. We thank Éric Meyer, Institut de Biologie de l’Ecole Normale Supérieure, Paris, for advice during set up of the *P. bursaria* RNAi approach.

## METHODS

### Culture conditions and media

In all RNAi experiments, *Paramecium bursaria* 186b (CCAP 1660/18) strain was used. For experiments requiring a non-photo-endosymbiotic *Paramecium* system for comparison, *Paramecium tetraurelia* nd7 strain was used. For eDicer analysis, *Paramecium bursaria* 186b and Yad1g1N strains were used^41,55^.

*Paramecium* cells were cultured in New Cereal Leaf – Prescott Liquid media (NCL). NCL media was prepared by adding 4.3 mgL^-1^ CaCl_2_.2H_2_O, 1.6 mgL^-1^ KCl, 5.1 mgL^-1^ K_2_HPO_4_, 2.8 mgL^-1^ MgSO_4_.7H_2_O to deionised water. 1 gL^-1^ wheat bran was added, and the solution boiled for 5 minutes. Once cooled, media was filtered once through Whatman Grade 1 filter paper and then through Whatman GF/C glass microfiber filter paper. Filtered NCL media was autoclaved at 121°C for 30 mins to sterilise prior to use.

NCL medium was bacterized with *Klebsiella pneumoniae* SMC and supplemented with 0.8 mgL^-1^ β-sitosterol prior to propagation. *Paramecium* cells were sub-cultured 1:9 into fresh bacterized NCL media once per month for *Paramecium bursaria* 186b, and once every two weeks for *Paramecium tetraurelia* nd7. *Paramecium* cultures were maintained at 18°C with a light-dark (LD) cycle of 12:12h.

### LysoTracker staining and fluorescent imaging

*Paramecium* cells were fixed using 0.5% paraformaldehyde and incubated for 20 mins at RT. Cells were then treated with 2 μM LysoTracker Green DND-26 (Invitrogen) to stain acidic vesicles such as the lysosome, and incubated in constant darkness for 2 hours at RT prior to imaging. Stained *Paramecium* cells were imaged on an ImageXpress Pico Automated Cell Imaging System at 10x magnification, using the Cy5 (absorbance – 630/40 nm, emission – 695/45 nm) and FITC (absorbance – 465/40 nm, emission – 525/30 nm) channels to capture algal chlorophyll autofluorescence and LysoTracker Green DND-26 fluorescence respectively.

### Gene synthesis and construct design

Sequences for plasmid constructs were synthesised *de novo* by either Genscript or SynBio Technologies, and cloned into an L4440 plasmid vector. Sequences and cloning sites for each plasmid construct are detailed in **Table S1**. All modified constructs were confirmed by Sanger sequencing (Eurofins Genomics).

### RNAi feeding

*Paramecium* was fed with *E. coli* transformed with an L4440 plasmid construct with paired IPTG-inducible T7 promoters, facilitating targeted gene knock-down through the delivery of complementary double-stranded RNA (dsRNA). L4440 plasmid constructs were transformed into *E. coli* HT115 competent cells and grown overnight on LB agar (50 μgmL^-1^ Ampicillin and 12.5 μgmL^-1^ Tetracycline) at 37°C. Positive transformants were picked and grown overnight in LB (50 μgmL^-1^ Ampicillin and 12.5 μgmL^-1^ Tetracycline) at 37°C with shaking (180 rpm). Overnight pre-cultures were back-diluted 1:25 into 50 mL of LB (50 μgmL^-1^ Ampicillin and 12.5 μgmL^-1^ Tetracycline) and incubated for a further 2 hours under the same conditions, until an OD_600_ of between 0.4 and 0.6 was reached. *E. coli* cultures were then supplemented with 0.4 mM IPTG to induce template expression within the L4440 plasmid, and incubated for a further 3 hours under the same conditions. *E. coli* cells were pelleted by centrifugation (3100 x *g* for 2 mins), washed with sterile NCL media, and pelleted once more. *E. coli* cells were then re-suspended in NCL media supplemented with 0.4 mM IPTG, 100 μgmL^-1^ Ampicillin, and 0.8 μgmL^-1^ β-sitosterol, and adjusted to a final OD_600_ of 0.1.

*Paramecium* cells were pelleted by gentle centrifugation in a 96-well plate (10 mins at 800 x *g*), taking care not to disturb the cell pellet by leaving 50 μl of supernatant, and re-suspended 1:4 into 200 μl of induced *E. coli* culture media (to make 250 μl total). Feeding was conducted daily for up to 14 days using freshly prepared bacterized media.

### Single-stranded construct design and confirmation

To create a version of L4440 which expressed only single-stranded (ssRNA), L4440 was digested with KpnI and PvuII (Promega), gel-purified (Wizard SV Gel and PCR Clean-Up System, Promega) and blunted using *PfuUltra* HF DNA polymerase (Agilent Technologies). The blunt vector was re-ligated using T4 DNA ligase (Thermo Scientific) and confirmed by sequencing (Eurofins Genomics), generating plasmid pDM004 which contains only a single T7 promoter. Fragments were then excised from their respective L4440 plasmids, or amplified by PCR (Q5 Polymerase; New England Biolabs) to swap the restriction sites, digested, and ligated into pDM004 to generate plasmids containing inserted fragments in sense [+] or antisense [-] orientation. Generated plasmids were transformed into HT115 *E. coli* for use in RNAi feeding experiments.

To confirm that these plasmid constructs generated only ssRNA, cultures of *E. coli* HT115 containing pDM005-1 ([-]ssRNA chimera) or pDM005-2 ([+]ssRNA chimera) were grown overnight at 37°C, 180 rpm in LB supplemented with 50 μgmL^-1^ ampicillin and 12.5 μgmL^-1^ tetracycline. Cultures were diluted 1:25 in fresh medium and grown to an OD_600_ of 0.4-0.6. Expression was then induced with 400 μM IPTG for 3 hrs, after which 1 mL of culture was pelleted by centrifugation for 2 mins at 3,100 x *g*. RNA was extracted using an RNeasy Mini Kit (Qiagen), following the manufacturer’s protocol for Total RNA Purification from Animal Cells. 20 μl of RNA was then treated with 10 μgmL^-1^ RNase A (Sigma-Aldrich) for 1 hour at 30°C in the presence of 300 mM NaCl (stabilising dsRNA and allowing RNase A to degrade only ssRNA^63^). A separate aliquot was left untreated, with 300 mM NaCl added to facilitate precipitation. All samples were then extracted with 1:1 phenol:chloroform and precipitated with 2 volumes of ethanol. Pellets were washed twice with 80% ethanol and re-suspended in nuclease-free water. RNA samples were then cleared of residual genomic DNA using the TURBO DNA-*free* Kit (Ambion), following the manufacturer’s protocol for routine DNase treatment. RT-PCR was then performed using the Qiagen OneStep RT-PCR kit following the manufacturer’s instructions, with 0.5 μL template RNA and 0.6 μM each primer (pDM005_RT_F: 5’-ACTTCAATGATTCGCAGCGG-3’ and pDM005_RT_R: 5’-AAGTAGCTGCTGTTCTCGGT-3’), generating an 85-nt PCR product. Cycling conditions were as detailed in the manufacturer’s protocol (30 cycles), with 1 min annealing at 50°C. PCR products were then resolved on a 2% agarose gel to assess for the presence/absence of amplification in each sample **(Figure S11).**

### qPCR analysis

RNA was extracted from *P. bursaria* 186b for gene expression analysis after three days of RNAi feeding. *Paramecium* cells (~10^3^ per culture) were pelleted by gentle centrifugation (800 x *g* for 10 mins), snap-frozen in liquid nitrogen, and stored at −80°C. RNA extraction was performed using TRIzol reagent (Invitrogen), following the manufacturer’s protocol after re-suspending each pellet in 900 μl TRIzol reagent. RNA was precipitated using GlycoBlue Co-precipitant (Invitrogen) to aid RNA pellet visualisation, and then cleared of residual DNA using the TURBO DNA-*free* Kit (Ambion), following the manufacturer’s protocol for routine DNase treatment.

RNA was reverse transcribed into single stranded cDNA using the SuperScript^®^ III First-Strand Synthesis SuperMix (Invitrogen), following the manufacturer’s protocol. Quantitative PCR (qPCR) was performed in a StepOnePlus Real-Time PCR system (Thermo Fisher Scientific). Reaction conditions were optimised using a gradient PCR, with a standard curve determined using 10-fold dilutions of *P. bursaria* cDNA: *EF-1α* (slope: −3.353; R^2^: 0.999; efficiency 98.740%), *HSP90* (slope: −3.319; R2: 0.998; efficiency 100.131%), *tub-β* (slope: – 3.378; R^2^: 0.992; efficiency 97.692%), and *actin* (slope: −3.349; R^2^: 0.983; efficiency 98.866%), using StepOne software v2.3. Each 20 μL reaction contained 10 μL PowerUp SYBR Green Master Mix (Thermo Fisher Scientific), 500 nM each primer (300 nM for *tub-β* and 1 μL (50 ng) cDNA. Each reaction was performed in duplicate for each of 3 biological replicates, alongside a ‘no-RT’ (i.e. non-reverse transcribed RNA) control to detect any genomic DNA contamination. Cycling conditions were as follows: UDG activation, 2 mins at 5O°C and DNA polymerase activation, 2 mins at 95°C, followed by 40 cycles of 15 secs, 95°C and 1 min at 55-65°C (*EF-1α* (58°C), *HSP90* (58°C), *tub-β* (57°C) and *actin* (58°C)). Primers pairs for each reaction are listed in **Table S5**. Each reaction was followed by melt-curve analysis, with a 60-95°C temperature gradient (0.3°C s^-1^), ensuring the presence of only a single amplicon, and ROX was used as a reference dye for calculation of C_T_ values. C_T_ values were then used to calculate the change in gene expression of the target gene in RNAi samples relative to control samples, using a derivation of the 2^-ΔΔCT^ algorithm^64^.

### sRNA isolation and sequencing

Total RNA for sRNA sequencing was extracted from *P. bursaria* (or free-living algal) cultures using TRIzol reagent (Invitrogen), as detailed above. To isolate sRNA from total RNA, samples were size separated on a denaturing 15% TBE-UREA polyacrylamide gel. Gels were prepared with a 15 mL mix with final concentrations of 15% Acrylamide/Bis (19:1), 8M UREA, TBE (89 mM Tris, 89 mM Borate, 2 mM EDTA), and the polymerisation started by the addition of 150 μL 10% APS (Sigma-Aldrich) and 20 μL TEMED (Sigma-Aldrich). Gels were pre-equilibrated by running for 15 mins (200 V, 30 mA) in TBE before RNA loading. The ladder mix consisted of 500 ng ssRNA ladder (50-1000nt, NEB#N0364S), and 5-10 ng of each 21 & 26-nt RNA oligo loaded per lane. The marker and samples were mixed with 2X RNA loading dye (NEB) and heat denatured at 90 °C for 3 mins before snap cooling on ice for 2 min prior to loading. Blank lanes were left between samples/replicates to prevent cross-contamination during band excision. Gels were then run for 50 mins (200V, 30 mA).

Once run, gels were stained by shaking (60 rpm) for 20 mins at RT in a 40 mL TBE solution containing 4 μL SYBR^®^ Gold Nucleic Acid Gel Stain. Bands of the desired size range (~15-30 nt) were visualised under blue light, excised and placed into a 0.5 mL tube pierced at the bottom by a 21-gauge needle, resting within a 1.5 mL tube, and centrifuged (16,000 x *g* for 1 min). 400 μL of RNA elution buffer (1M Sodium acetate pH 5.5 and 1mM EDTA) was added to the 1.5 mL tube containing centrifuged gel slurry, and the empty 0.5 mL tube discarded. Gel slurry was manually homogenized until dissolved using a 1 mL sterile plunger and incubated at RT for 2 hours with shaking at 1,400 rpm.

Solutions containing RNA elution buffer and gel slurry were transferred to a Costar Spin-X 0.22 μm filter column and centrifuged (16,000 x *g* for 1 min). The filter insert containing acrylamide was discarded. 1 mL of 100% EtOH was added to each solution, alongside 15 μg of GlycoBlue^™^ Coprecipitant (Invitrogen) to aid sRNA pellet visualisation, and stored overnight at −80°C to precipitate. Precipitated solutions were centrifuged at 4°C (12,000 x *g* for 30 mins), and the supernatant discarded. sRNA pellets were washed with 500 μL of cold 70% EtOH (12,000 x *g* for 15 mins at 4°C), and air dried in a sterile PCR hood for 10 mins, before re-suspending in 15 μL of RNAse-free water and storage at −80°C.

### sRNA-seq and read processing

sRNA concentrations were determined using an Agilent 2100 Bioanalyzer, following the Agilent Small RNA kit protocol, and all samples matched to 0.7 ngmL^-1^ prior to sequencing. Library preparation and subsequent RNA-seq was performed for 54 samples using 50-bp paired-end, rapid run across four lanes on an Illumina HiSeq 2500, yielding ~120-150 million paired-end reads per lane (~9-11 million paired-end reads per sample).

The raw paired-end reads from the RNA-seq libraries were trimmed using Trim Galore in order to remove barcodes (4-nt from each 3’- and 5’-end) and sRNA adaptors, with additional settings of a phred-score quality threshold of 20 and minimum length of 16-nt. Result were subsequently checked with FastQC.

### Assigning sRNAs to the algal endosymbiont transcript bins

Trimmed reads were mapped against the ‘endosymbiont’ dataset of assembled transcripts using the HISAT2 alignment program with default settings. Post-mapping, the BAM files were processed using SAMTOOLS and a set of custom scripts (https://github.com/guyleonard/paramecium) to produce a table of mapped read accessions and their respective read lengths. Using these exported count tables, of mapped reads per read length, transcripts with >10 hits for read lengths between 21-25 nt were searched using reciprocal BLASTX against the NCBI non-redundant ‘nr’ proteins sequence database, in order to assign taxonomic identity to each transcript. This allowed filtering of the main ‘endosymbiont’ dataset into subsets corresponding to either: algal cytoplasmic mRNA, algal cytoplasmic rRNA, algal plastid RNA or algal mitochondrial RNA; host RNA contamination; bacterial RNA contamination; or vector RNA contamination. All identified algal cytoplasmic rRNA (28 transcripts), plastid RNA (56 transcripts) or mitochondrial RNA (18 transcripts) sequences were sorted into new datasets representing each RNA species. Host (5 transcripts) and bacterial (34 transcripts) contamination were sorted into respective ‘host’ or ‘bacterial’ datasets. Vector (1 transcript) and all unidentifiable sequences (20 transcripts) were separated into a dataset labelled as ‘other’. All remaining transcripts with >10 sRNA hits for read sizes 21-25 nt in the original dataset now represented putative algal mRNA transcripts (i.e. for 23-nt reads this represented 3,659 sRNA reads mapping to 148 transcripts). In this algal mRNA dataset, we also included transcripts that fell below the >10 sRNA hit threshold for manual curation, and which therefore may correspond to reads mapping to all of the above bins (i.e. for 23-nt reads this represented 1,949 reads mapping to 605 transcripts in total, of which 1,408 reads were mapping to 468 transcripts which were not manually curated, so could be affected by contamination). This manual binning process was carried out in order to double check the original automated binning post transcriptome assembly, and to check for possible chimeric host-algal transcript sequences produced as a by-product of cDNA synthesis and transcriptome sequencing and assembly. This process collectively allowed accurate segregation of the existing ‘endosymbiont’ transcript bin into discrete subsets based on RNA ‘species’.

Using the HISAT2 alignment program with default settings, trimmed reads were mapped once more against the newly filtered algal mRNA, rRNA, plastid and mitochondrial datasets from above. Count tables of mapped reads per read length were once again generated from the BAM files, and used to plot a size distribution of 21-29 nt endosymbiont derived sRNA abundance per RNA species. Size distributions of sRNA abundance for each sample were plotted using the R programming language packages; tidyverse, grid.extra and ggplot2 in R Studio.

### eDicer methods for identifying putative RNA-RNA interactions

To predict putative mRNA-mRNA interactions using eDicer (https://github.com/fmaguire/eDicer), both the ‘host’ and ‘endosymbiont’ transcript bins processed from transcriptome data for *P. bursaria* Yad1g1N^55^ were further filtered to minimise the risk of false positives for host and endosymbiont cross-comparisons. For the ‘host’ bins, any transcript with >90% ID BLASTN hit to the following genome assemblies were removed: *Chlamydomonas reinhardtii* cc503 cw92 mt, *Chlamydomonas reinhardtii* v3, *Chlorella sorokiniana* 1228 v2, *Chlorella sorokiniana* DOE1412 v3, or *Chlorella sorokiniana* utex1230 lanl v2 assemblies from Los Alamos National Labs Greenhouse genome database. Similarly, for the ‘endosymbiont’ bins, any transcript with >90% ID BLASTN hit to the following ParameciumDB genome assemblies were removed: *P. biaurelia* V1-4 v1, *P. caudatum* 43c3d v1, *P. bursaria* MAC 110224 v1, *P. decaurelia* MAC 223 v1, *P. dodecaurelia* MAC 274 v1, *P. jenningsi* MAC M v1, *P. novaurelia* MAC TE v1, *P. octaurelia* K8 CA1, *P. quadecaurelia* MAC NiA v1, *P. primaurelia* Ir4-2 v1, *P. tetraurelia* MAC 51 with and without IES, or *P. tetraurelia* MAC. Additionally, the reference (refseq) genome of *Escherichia coli* MG1655 (NCBI acc.: NZ_CP012868.1) and *Klebsiella pneumoniae* HS11286 (NCBI acc.: NC_016845.1) were used as ‘food’ comparison datasets.

The filtered ‘endosymbiont’ transcript bin, *E. coli* CDS and *K. pneumoniae* CDS were decomposed into all possible 21-23 nt reads with jellyfish v2.2.10 and aligned with 95% identity to the ‘host’ bin using Bowtie v1.2.3 (recommended for short alignments). These *in*-silico RNA-RNA interaction simulations were performed using a wrapper tool we created named eDicer (v1.0.0) (https://github.com/fmaguire/eDicer). The number of distinct aligning k-mers (i.e. putative interactions of unique sequences) for each pair of bins were then normalised by dividing by the total number of distinct k-mers in both bins, and x 100, to calculate a Jaccard Index % (i.e. normalised set similarity).

To predict putative ‘lethal’ RNA-RNA interactions, a separate analysis was conducted using a subset of each dataset that was putatively homologous to a yeast ‘lethal’ gene database^56^. This ‘lethal’ dataset contained genes known to be conditionally essential in *Saccharomyces cerevisiae*. These putative ‘lethal’ homologues for each dataset were identified using a tBLASTx search of the yeast ‘lethal’ database^56^ with a gathering threshold set at 1e-10 with a minimum of 50% sequence identity. Once curated, each ‘lethal’ dataset was subject to eDicer analysis as described above.

To predict putative rRNA-rRNA interactions using eDicer, a further analysis was conducted using a dataset consisting of full-length ribosomal RNA (rRNA) clusters for *Microactinium conductrix* [NCBI acc.: ASM224581v2]^65^, *K. pneumoniae* [NCBI acc.: NC_016845.1]^66^, *E. coli* [NCBI acc.: NZ_CP012868.1], *P. bursaria* Yad1g1N^55^ and *P. bursaria* 186b. Once again, each dataset was subject to eDicer analysis as described above. All *in-silico* RNA-RNA interaction predictions were plotted using the R programming language packages; tidyverse, grid.extra and ggplot2 in R Studio.

### Manual curation of additional host transcript bins

In order to identify host transcripts that could be impacted by putative RNA-RNA interactions as a result of endosymbiont derived sRNA exposure, trimmed Illumina reads for all sRNA sequencing samples were mapped to the ‘endosymbiont mRNA’ transcript dataset. Resulting mapping files were filtered with custom scripts (https://github.com/guyleonard/paramecium) producing tables of mapped hits with their respective read lengths. Reads from all tables, of length 23-nt only, were then extracted with SEQTK (resulting in 3,690 total). A BLASTn search of these 23-nt endosymbiont reads was conducted against the ‘host’ transcript dataset, to identify host transcripts with ≥ 95% identity over a 23-nt region. This resulted in three candidate host transcripts. These three host transcripts were searched using BLASTx against the NCBI non-redundant ‘nr’ protein database, resulting in one transcript with 100% sequence identity to both endosymbiont and host HSP90. For the remaining two transcripts, SNPs present in the identified 23-nt mapped sequence indicate that these are putative host sequences with 2-nt of mismatch compared to the respective endosymbiont sequence, and were therefore unlikely to represent a product of RNA-RNA interactions between endosymbiont and host as a result of endosymbiont derived 23-nt sRNA exposure.

In order to identify host transcripts that could be classified as ‘non-hit’ transcripts (i.e. host transcripts sharing a low level of sequence identity with algal transcripts over 23-nt regions), the ‘host’ dataset was searched using an organism specific BLASTn (Chlorellaceae [NCBI: taxid35461]) against the NCBI nucleotide ‘nr/nt’ database with a minimum expectation of 1e-05. Any host transcripts sharing >20-nt sequence identity with algal transcripts over a 23-nt region were rejected. This process was repeated until a dataset of 20 ‘non-hit-algal’ host transcripts were identified. These putative ‘non-hit-algal’ host transcripts were then searched using BLASTx against the NCBI non-redundant ‘nr’ protein database, to confirm that these 20 transcripts were derived from *Paramecium* and were not bacterial contamination.

## DATA AND SOFTWARE AVAILABILITY

The raw reads generated during sRNA sequencing are available on the NCBI Sequence Read Archive (accessions: SAMN14932981, SAMN14932982). All other datasets are available on Figshare (https://doi.org/10.6084/m9.figshare.c.4978160.v1), under the relevant headings. Custom scripts for sRNA read processing (https://github.com/guyleonard/paramecium, https://doi.org/10.5281/zenodo.4638888) and eDicer comparative analysis (https://github.com/fmaguire/eDicer, https://doi.org/10.5281/zenodo.4659378) are available on GitHub and archived within the Zenodo repository.

