## Supplemental Material for "Emergent RNA-RNA interactions can promote stability in a nascent phototrophic endosymbiosis"

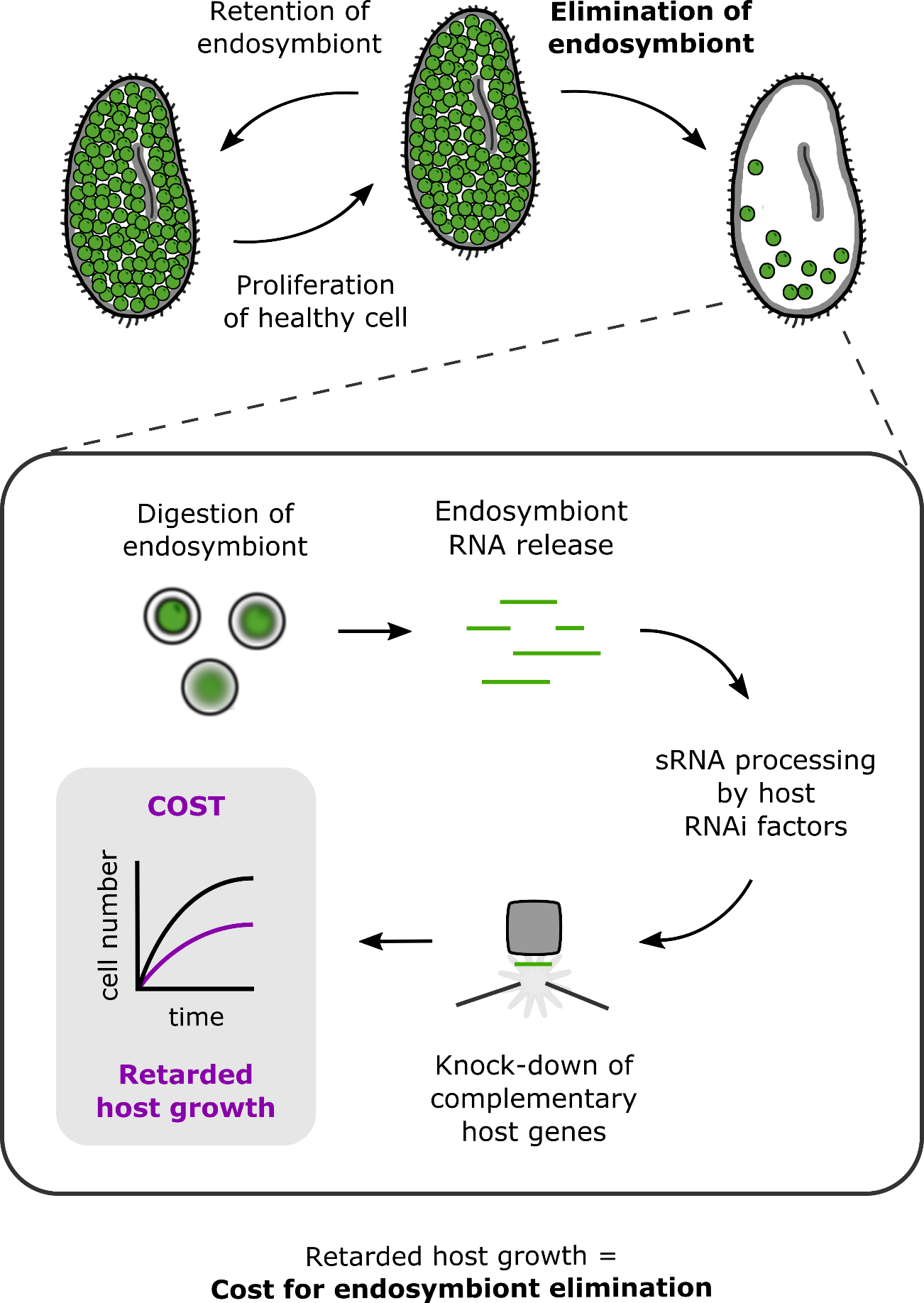

**Figure S1. An RNAi-mediated physiological cost for endosymbiont digestion.** Graphical abstract demonstrating how endosymbiont derived RNA released during endosymbiont digestion is processed by the host RNAi system, resulting in knock-down of complementary host genes which imposes a cost to host growth. We propose this mechanism to be mediated by host Dicer, AGO-Piwi, Pds1 and RdRP proteins. Such a cost would effectively ‘punish’ the host for digestion of the endosymbiont population, generating an evolutionary outcome in which maintenance of the interaction is favoured over breakdown. Consequently, this mechanism would promote stability and act to drive cooperation within the nascent endosymbiotic interaction. We term this process of host gene knock-down in response to endosymbiont RNA processing by host RNAi factors: ‘RNAi-collisions’.

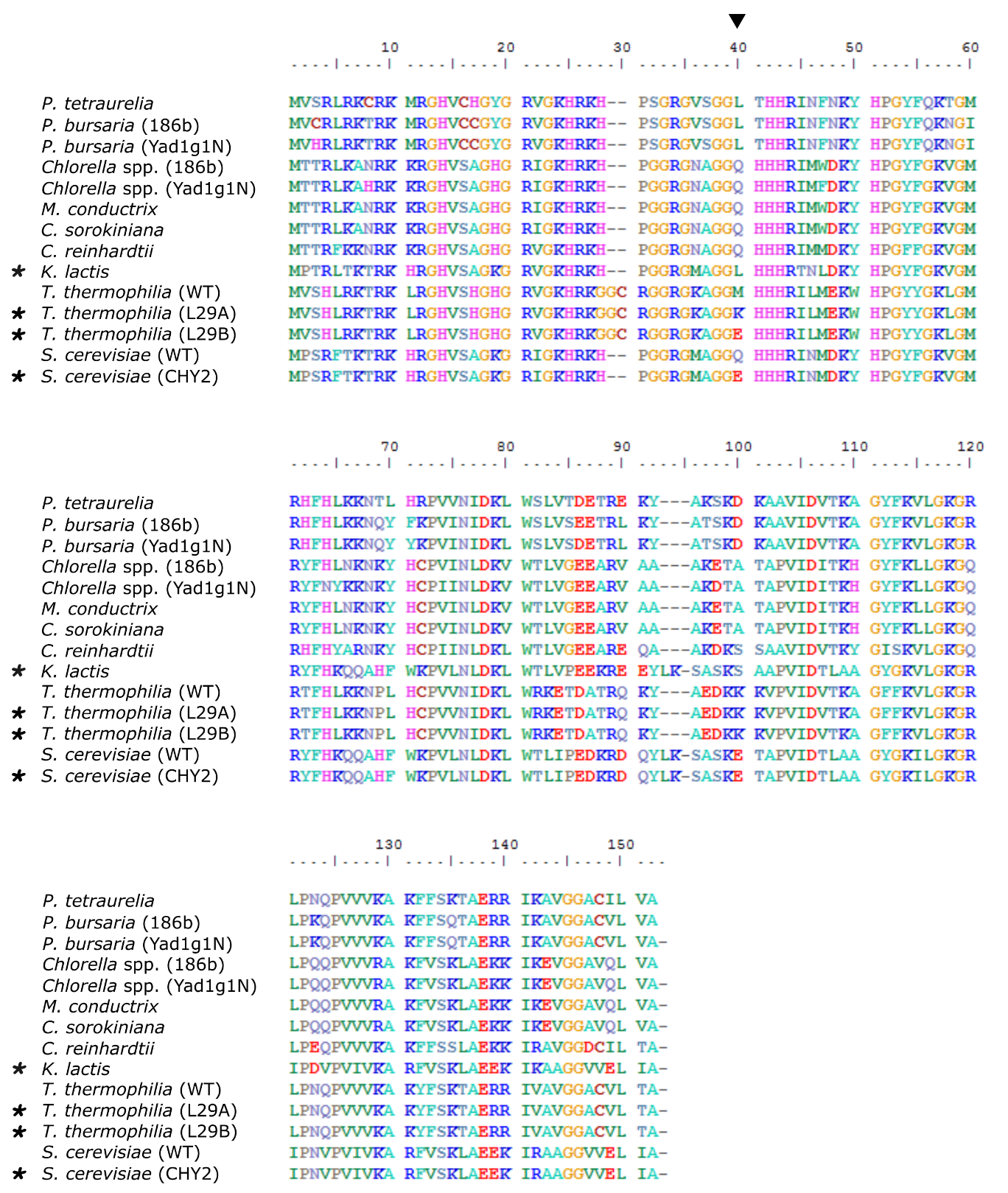

**Figure S2. *Paramecium* encodes an amino acid residue change that confers cycloheximide resistance, while the algal endosymbionts do not.** Alignment of RPL29A from ciliate, algal and fungal amino acid sequences, and from *P. bursaria* transcriptome predicted protein data. Asterisks denote known cycloheximide resistant species. Cycloheximide resistance in *T. thermophilia (L29A* or *L29B)* and *S. cerevisiae (CHY2)* was conferred through a single point mutation^1^ resulting in an amino acid residue change of ‘Met40’ (black arrow; residue 38 in *P. bursaria* RPL29A). Amino acid alignment data for RPL29A is available on Figshare ([10.6084/m9.figshare.12301973](https://doi.org/10.6084/m9.figshare.12301973)).

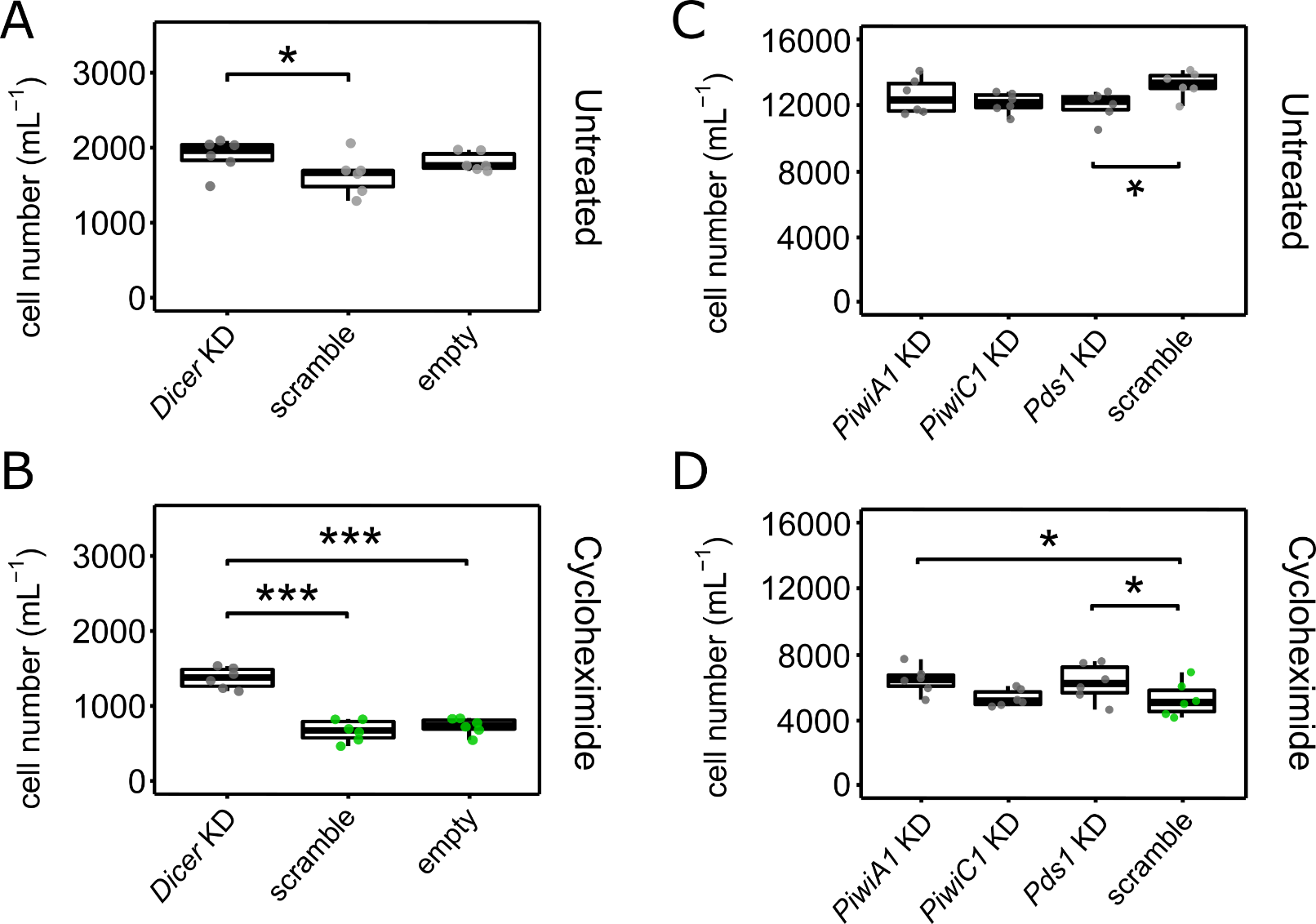

**Figure S3. Exposure to *Dicer*, *PiwiA1* or *Pds1* dsRNA significantly rescues *P. bursaria* culture growth during treatment with cycloheximide to induce endosymbiont digestion.** (**A**-**B**) *P. bursaria* cell number after 8 days of treatment with cycloheximide at 50 µgmL^-1^, compared to untreated control cultures. *P. bursaria* cultures were simultaneously fed for 12 days (starting four days prior to drug treatment) with *E. coli* expressing; *Dicer* dsRNA, non-hit ‘scramble’ dsRNA, or an empty vector control. (**C**-**D**) *P. bursaria* cell number after 6 days of treatment with cycloheximide at 50 µgmL^-1^, compared to untreated control cultures. *P. bursaria* cultures were fed for 18 days (starting 12 days prior to treatment) with *E. coli* expressing; *PiwiA1* dsRNA, *PiwiC1* dsRNA, *Pds1* dsRNA, or a non-hit ‘scramble’ dsRNA control. All boxplot data are represented as max, upper quartile (Q3), mean, lower quartile (Q1) and min values of six biological replicates. Significance calculated as *p ≤ 0.05, ***p ≤ 0.001 using a generalized linear model with quasi-Poisson distribution. This figure is an extension of **Figure 1D** & **E**, showing the raw count data used to calculate the % change in cell number upon treatment with cycloheximide.

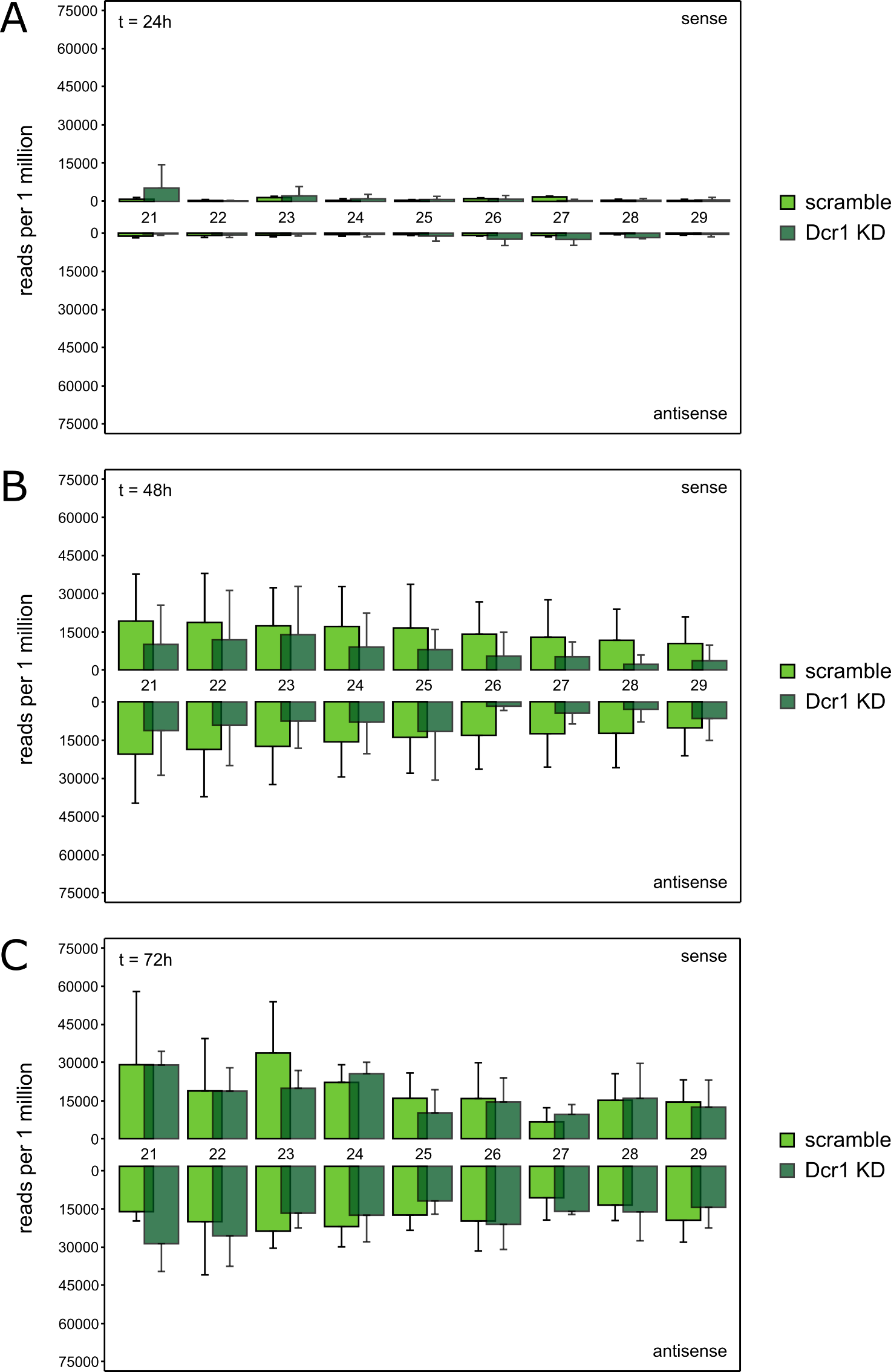

**Figure S4. Endosymbiont mRNA-derived sRNA abundance increases during cycloheximide treatment, dependent on *Dicer* processing.** Size distribution of sRNA mapped to endosymbiont cytoplasmic mRNA transcripts. sRNA was extracted from *P. bursaria* cultures on day 1, 2 and 3 of cycloheximide treatment (50 µgmL^-1^), or from an untreated control. *P. bursaria* cultures were fed with HT115 *E. coli* transformed to express ‘scramble’ or Dicer dsRNA. Feeding was conducted daily for four days prior to cycloheximide treatment, and continued throughout. Data are represented as mean ± SD of three biological replicates, and normalised to total endosymbiont mRNA-mapping 21-29 nt reads across all datasets. Curated ‘endosymbiont’ transcript bins used for sRNA mapping are available on Figshare ([10.6084/m9.figshare.12301736](https://doi.org/10.6084/m9.figshare.12301736)). This figure relates to **Figure 2**, and demonstrates that an increased abundance of reads mapping to endosymbiont-derived cytoplasmic mRNA is observed in response to 2-3 days of cycloheximide treatment.

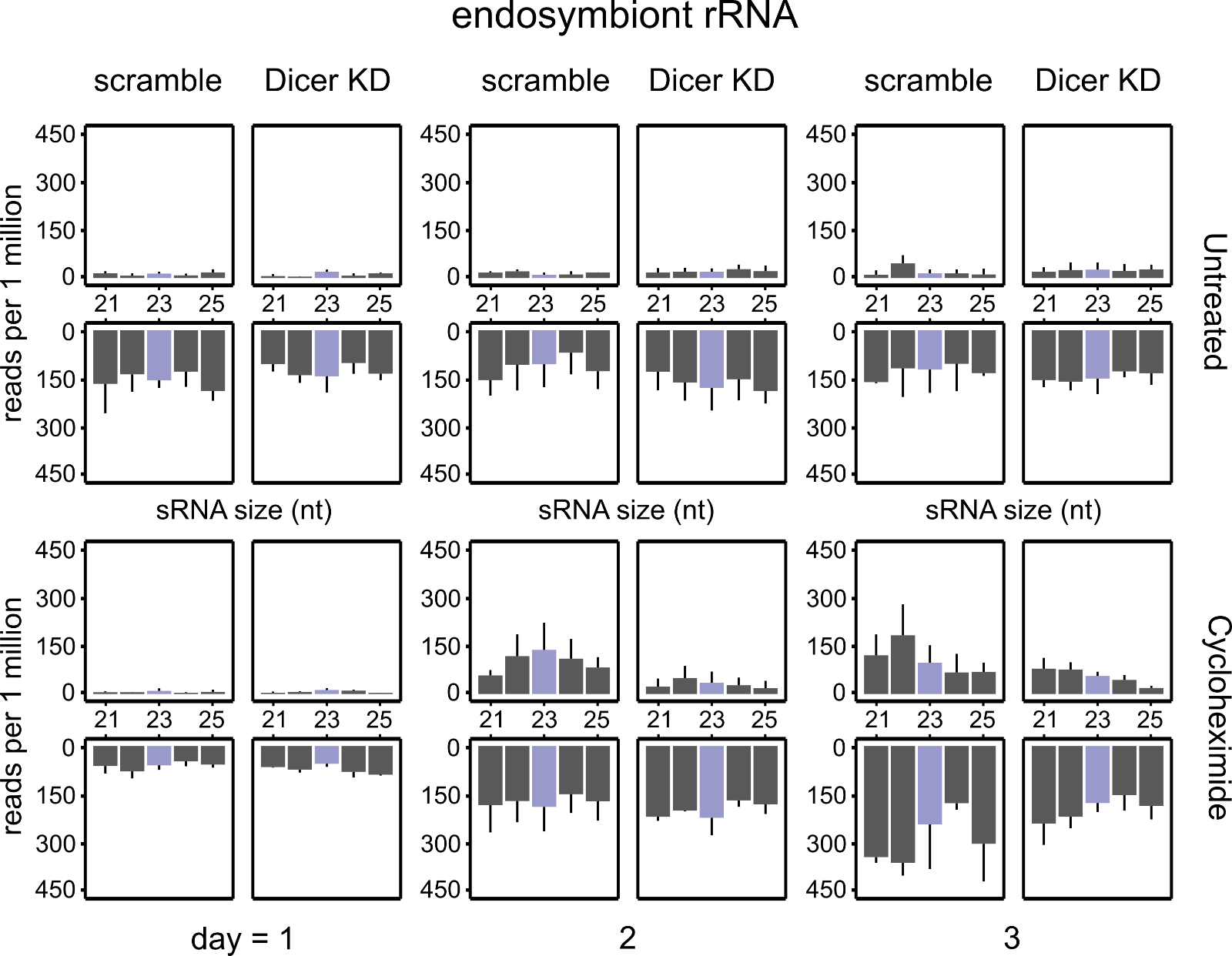

**Figure S5. Endosymbiont rRNA-derived sRNA abundance also increases during cycloheximide treatment, however background processing of rRNA occurs independent of treatment.** Size distribution of sRNA mapped to endosymbiont cytoplasmic rRNA transcripts. sRNA was extracted from *P. bursaria* cultures on day 1, 2 and 3 of cycloheximide treatment (50 µgmL^-1^), or from an untreated control. *P. bursaria* cultures were fed with HT115 *E. coli* transformed to express ‘scramble’ or Dicer dsRNA. Feeding was conducted daily for four days prior to cycloheximide treatment, and continued throughout. Data are represented as mean ± SD of three biological replicates, and normalised to total 21-25 nt reads per sample. Curated ‘endosymbiont’ transcript bins used for sRNA mapping are available on Figshare ([10.6084/m9.figshare.12301736](https://doi.org/10.6084/m9.figshare.12301736)). This figure relates to **Figure 2**, and demonstrates that an increased abundance of reads mapping to endosymbiont-derived cytoplasmic rRNA is also observed in response to 2-3 days of cycloheximide treatment. In addition, this figure demonstrates the occurrence of antisense oriented sRNA in both untreated and cycloheximide treated conditions, representing putative background processing of endosymbiont derived rRNA independent of cycloheximide treatment.

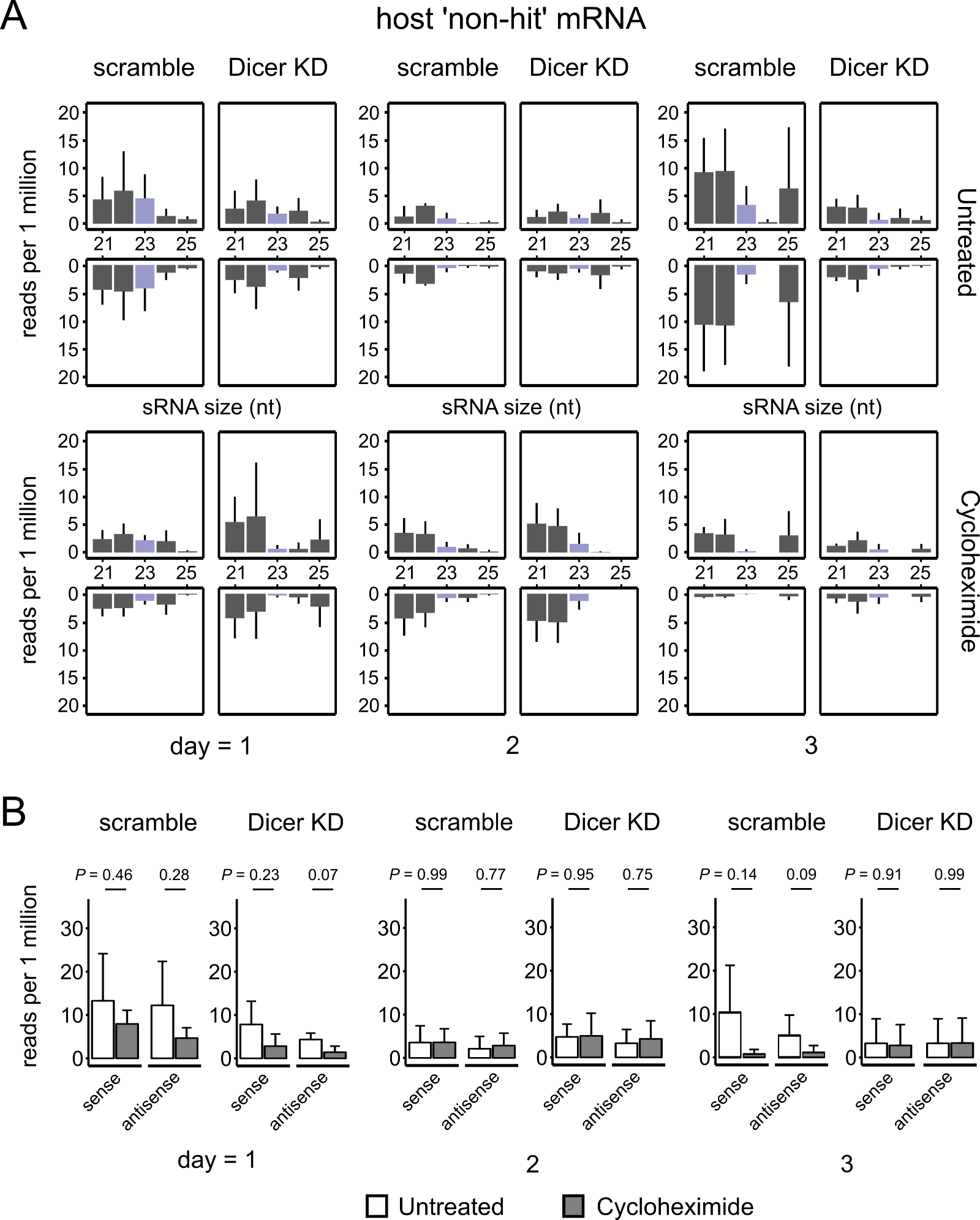

**Figure S6. Host sRNA abundance does not significantly increase during cycloheximide treatment, for host transcripts sharing low sequence identity with endosymbiont RNA.** (**A**) Size distribution of sRNA mapped to 20 ‘non-hit’ host mRNA transcripts sharing <87% sequence identity with algal mRNA over 23-nt regions, to ensure that these host transcripts were unaffected by putative RNA-RNA interactions derived from the endosymbiont. sRNA was extracted from *P. bursaria* cultures on day 1, 2 and 3 of cycloheximide treatment (50 µgmL^-1^), or from an untreated control at the same time points. *P. bursaria* cultures were fed with HT115 *E. coli* transformed to express ‘scramble’ or Dicer dsRNA. Feeding was conducted daily for four days prior to cycloheximide treatment, and continued throughout. Data are represented as mean ± SD of three biological replicates, and normalised to total 21-25 nt reads per sample. (**B**) Comparison of 23-nt reads mapping to 20 ‘non-hit’ host transcripts in untreated or cycloheximide treated *P. bursaria* cultures. Values are presented on a log scale to aid visualisation. Data are represented as mean ± SD of three biological replicates, and normalised to total 23-nt reads per sample. *P* values were calculated using a paired t-test and are displayed within the figure. Curated ‘host’ transcript bins used for sRNA mapping are available on Figshare ([10.6084/m9.figshare.12301736](https://doi.org/10.6084/m9.figshare.12301736)). This figure relates to **Figure 2**, and demonstrates that for host mRNA transcripts which do not share a high level of sequence identity with endosymbiont RNA, sRNA abundance does not increase significantly during cycloheximide treatment.

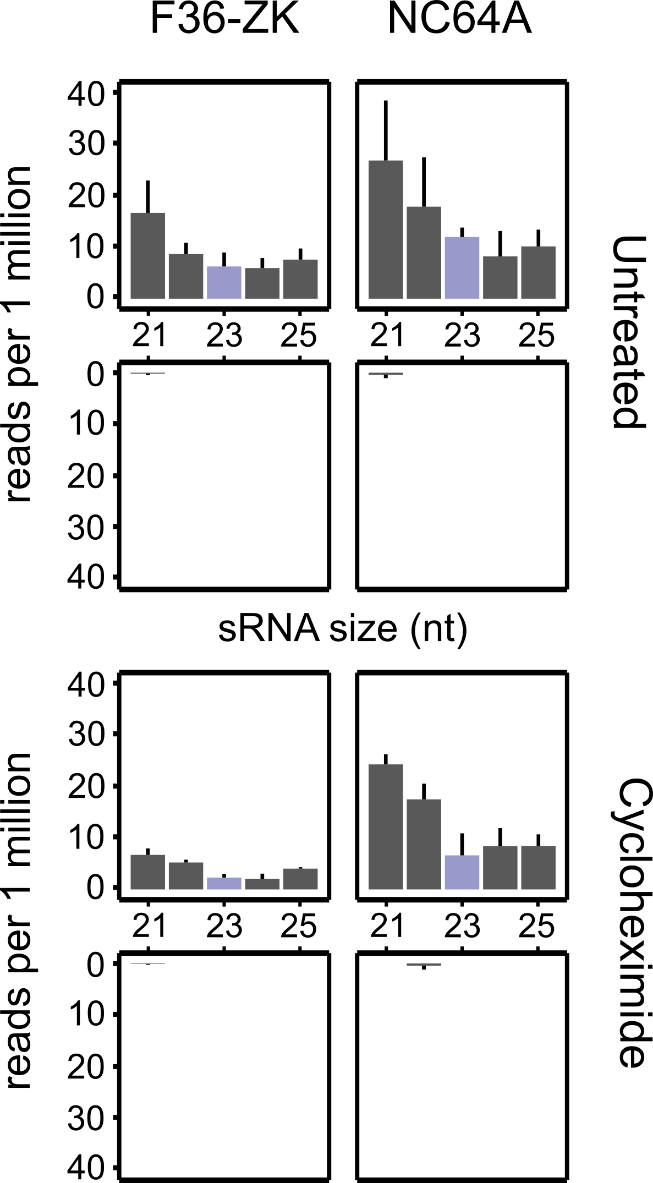

**Figure S7. Free-living algal sRNA abundance does not increase during cycloheximide treatment.** (**A**) Size distribution of sRNA mapped to endosymbiont mRNA. sRNA was extracted from free-living algal cultures (*Chlorella variabilis* F36-ZK and *Chlorella variabilis* NC64A, both known endosymbionts of *P. bursaria*) after 4 days of cycloheximide treatment (50 µgmL^-1^), or from an untreated control. Data are represented as mean ± SD of three biological replicates, and normalised to total 21-25 nt reads per sample. Curated ‘endosymbiont’ transcript bins used for sRNA mapping are available on Figshare ([10.6084/m9.figshare.12301736](https://doi.org/10.6084/m9.figshare.12301736)). This figure relates to **Figure 2**, and demonstrates that an increase in algal sRNA abundance upon treatment with cycloheximide is likely a result of host RNA processing within the *P. bursaria*-*Chlorella* spp. endosymbiotic system.

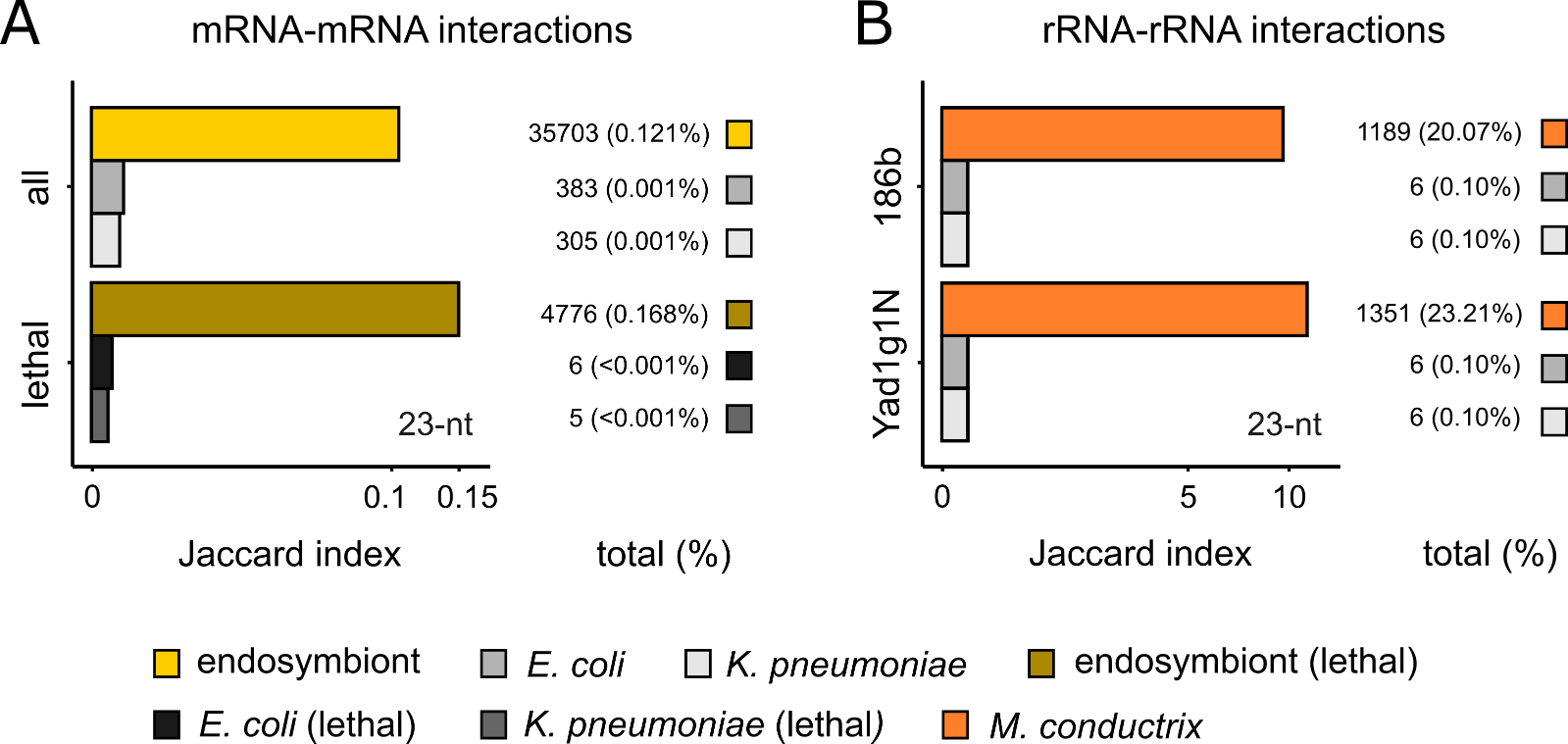

**Figure S8. Comparison of Transcriptome Data Reveals the Potential for Host Transcript Interaction by Endosymbiont Derived RNAs. (A)** Jaccard (% similarity) Index demonstrating the proportion of endosymbiont (yellow), *E. coli* (grey) or *K. pneumoniae* (light grey) mRNA transcripts that map with >91% sequence identity to host mRNA transcripts over a 23-nt region in *P. bursaria* (Yad1g1N). These regions of 23-nt overlap represent putative mRNA-mRNA interactions between endosymbiont / bacterial-food and host RNA. Also shown are the proportion of endosymbiont (dark yellow), *E. coli* (black) or *K. pneumoniae* (dark grey) mRNA transcripts that map with >91% sequence identity to putatively ‘lethal’ host mRNA transcripts (transcripts possessing a high sequence identity to the yeast conditionally essential gene database (Cotton and McInerney, 2010) over a 23-nt region in *P. bursaria* (Yad1g1N). Using these datasets, we identified an approximately 120-fold (total) to 170-fold (‘lethal’) increase in the number of putative 23-nt mRNA-mRNA interactions predicted between endosymbiont-and-host compared to between bacterial food-and-host RNA. **(B)** Jaccard (% similarity) Index demonstrating the proportion of *M. conductrix* (orange), *E. coli* (dark grey) or *K. pneumoniae* (light grey) rRNA transcripts that map with >91% sequence identity to host rRNA transcripts over a 23-nt region in *P. bursaria* (186b and Yad1g1N). These regions of 23-nt overlap represent putative rRNA-rRNA interaction between endosymbiont / bacterial-food and host RNA. Using this dataset, consisting of full-length ribosomal RNA (rRNA) clusters, we identified an approximately 200-fold (186b) to 230-fold (Yad1g1N) increase in the number of putative 23-nt rRNA-rRNA interactions predicted between endosymbiont-and-host compared to between bacterial food-and-host. (**A-B**) For all panels, Jaccard Index values are presented on a square root scale to aid visualisation. Values displayed on the right indicate the number of distinct host 23-nt regions identified as being hit by the respective endosymbiont (or model food bacterium) 23-nt dataset. Values in brackets indicate the percentage of total host 23-nt k-mers that these 23-nt hits represent. Curated ‘host’ and ‘endosymbiont’ transcript bins used for eDicer comparative analyses are available on Figshare ([10.6084/m9.figshare.12302597](https://doi.org/10.6084/m9.figshare.12302597)). See also **Table S2** for data analysis relating to all predicted RNA-RNA interactions identified using eDicer. For a full overview of the eDicer process, see **Supplementary Methods**.

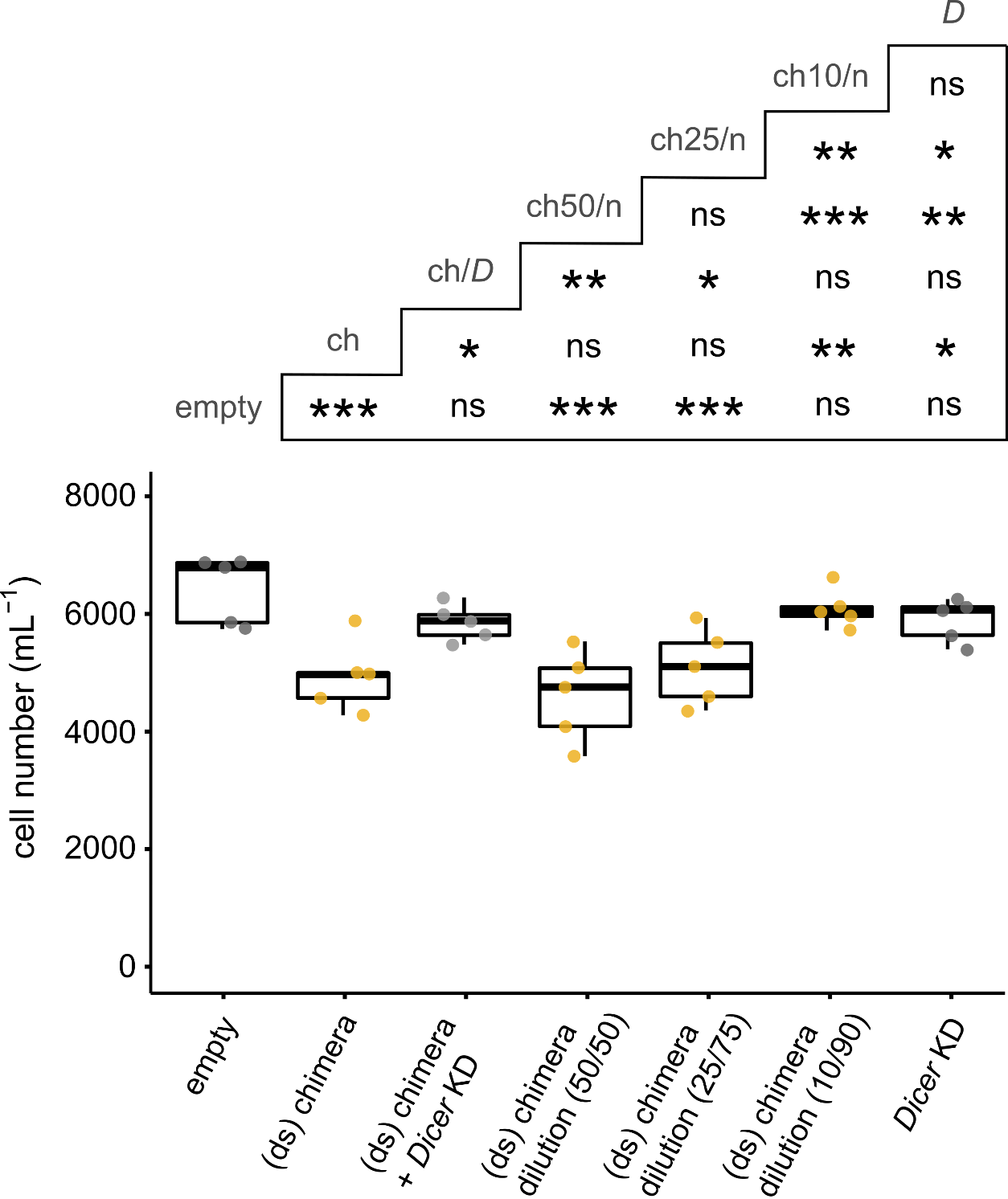

**Figure S9. Exposure to a synthetic endosymbiont derived chimera dsRNA diluted with ‘non-hit’ nonsense dsRNA also results in culture growth retardation in *P. bursaria*.** *P. bursaria* cell number after 12 days of feeding with *E. coli* expressing; chimera dsRNA or chimera dsRNA mixed with nonsense dsRNA (yellow); chimera dsRNA mixed with *Dicer* dsRNA (grey; rescue); *Dicer* dsRNA (grey); or an empty vector control (white). (ds, double-stranded). Multiple vector delivery of chimera/nonsense was conducted at a 50:50, 25:75 or 10:90 ratio of *E. coli* strain exposure during feeding. Multiple vector delivery of chimera/Dicer was conducted at a 50:50 ratio. Boxplot data are represented as max, upper quartile (Q3), mean, lower quartile (Q1) and min values of six biological replicates. Asterisks displayed in the grid above denote pairwise significance values, calculated as *p ≤ 0.05, **p ≤ 0.01, ***p ≤ 0.001, ‘ns’ no significance, using a generalized linear model with quasi-Poisson distribution. This data is an extension of **Fig. 3C**, and demonstrates that a relative dilution of endosymbiont-derived chimera dsRNA can also facilitate a cost to host growth.

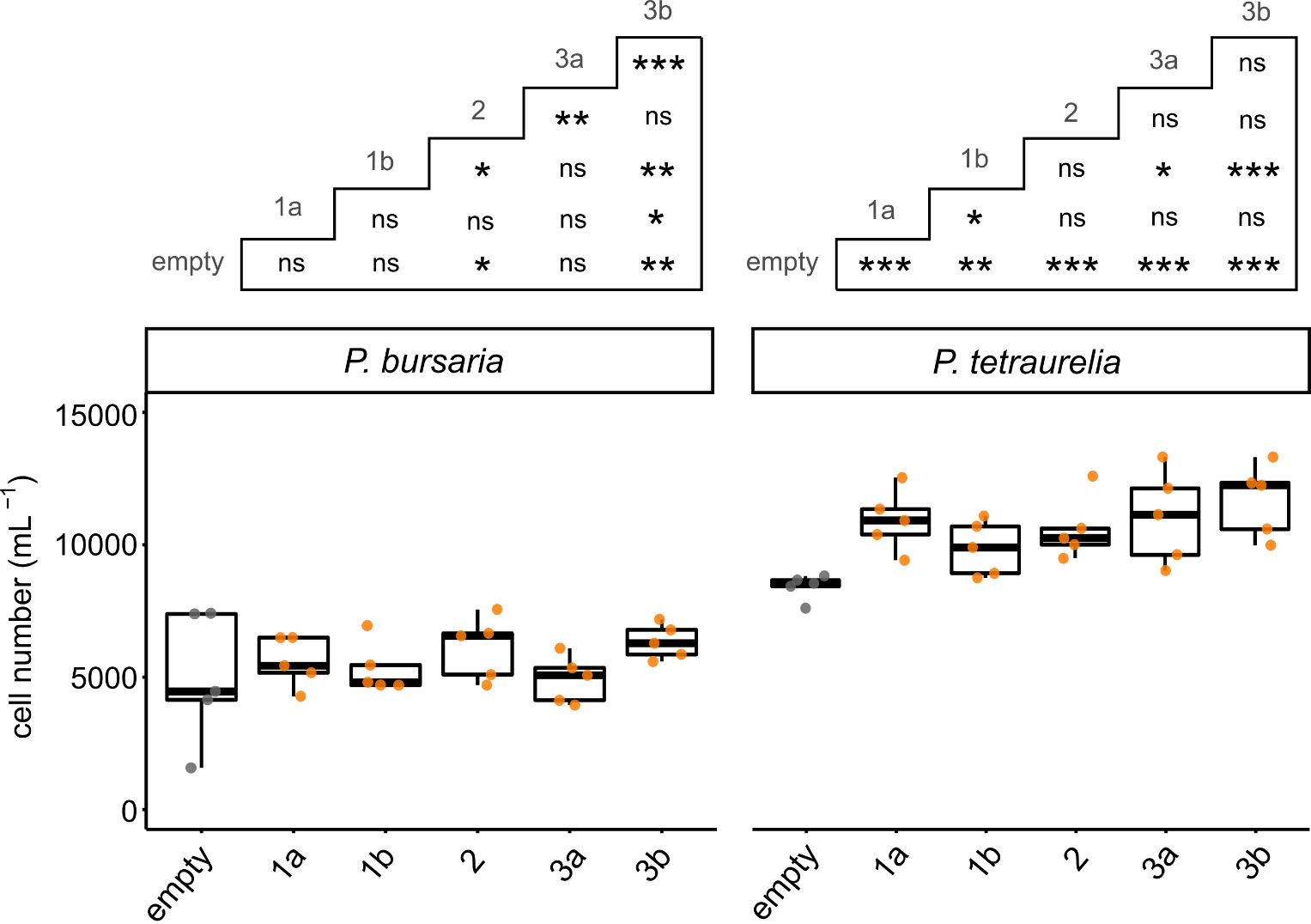

**Figure S10. Exposure to SSU rRNA derived dsRNA does not result in culture growth retardation in *Paramecium*.** *Paramecium* cell number in *P. bursaria* or *P. tetraurelia* cultures after 12 days of feeding with *E. coli* expressing rRNA interaction fragment dsRNA (orange; **Table S4**); or an empty vector control (white). Identified rRNA interaction fragments were comprised of 71-109 nt of small subunit (SSU) rRNA sharing multiple regions of >90% sequence identity between endosymbiont and host, and were derived from either the host (1a/2/3a) or endosymbiont (1b/2/3b) with 100% respective identity. Upon exposure to rRNA derived dsRNA, no significant retardation to culture growth was observed in either *P. bursaria* or *P. tetraurelia.* Boxplot data are represented as max, upper quartile (Q3), mean, lower quartile (Q1) and min values of five biological replicates. Asterisks displayed in the grid above denote pairwise significance values (right: down), calculated as *p ≤ 0.05, **p ≤ 0.01, ***p ≤ 0.001, ‘ns’ no significance, using a generalized linear model with quasi-Poisson distribution. These data demonstrate that either the host rRNA is effectively shielded from RNAi mediated interactions; that the rRNA population is too abundant for a significant effect to become apparent via induction of RNAi; or that these rRNA fragments are not efficiently processed by the *Paramecium* RNAi machinery.

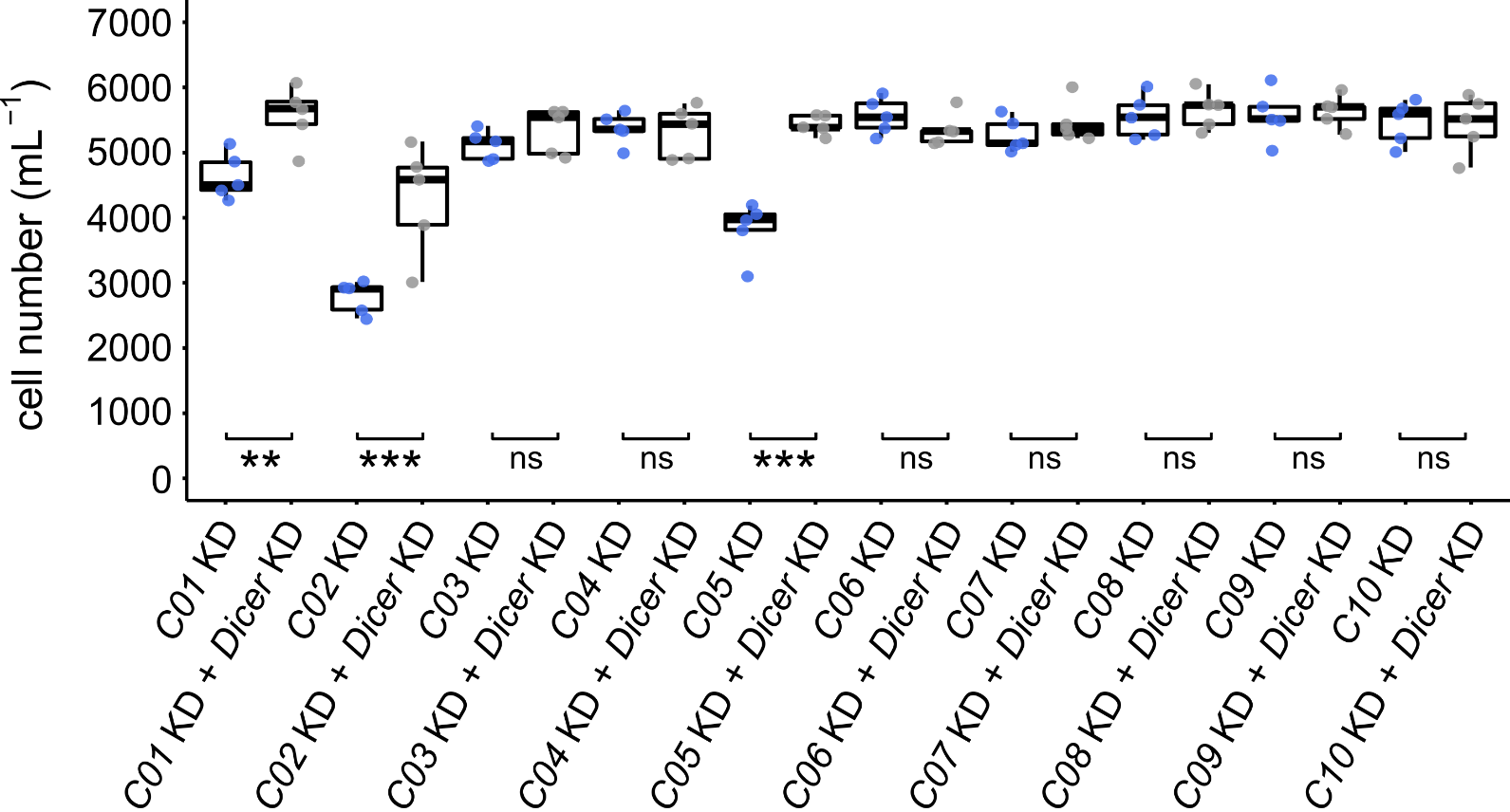

**Figure S11. Individual knock-down of the ten host targets corresponding to each constituent mRNA interaction fragment of the synthetic endosymbiont derived chimera, using 450-nt *P. bursaria* templates to target the wider host gene directly.** *P. bursaria* cell number after 12 days of feeding with *E. coli* expressing 450-nt dsRNA corresponding to host homologues of each of the ten-constituent endosymbiont derived chimera mRNA interaction fragments (yellow), compared to *Dicer* dsRNA mixed controls (grey; rescue phenotype). Multiple vector delivery was conducted at a 50:50 ratio during feeding. C01, C02 and C05 correspond to host genes: *EF1-α*, *HSP90* and *tub-β*. Significance calculated as ***p ≤ 0.001, ‘ns’ no significance, using a generalized linear model with quasi-Poisson distribution. This figure is an extension of **Figure 3D**, demonstrating which constituent regions of the synthetic endosymbiont derived chimera result in a cost to host growth in *P. bursaria* when the wider corresponding host gene is targeted directly.

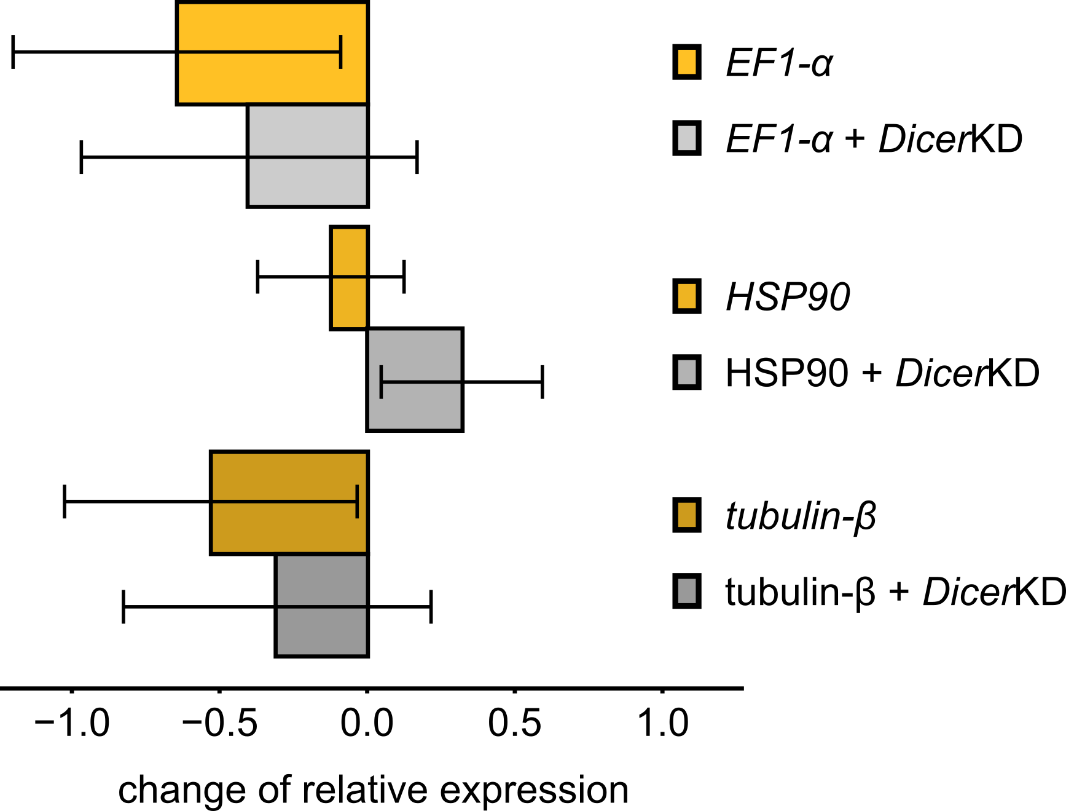

**Figure S12. Delivery of synthetic endosymbiont chimera dsRNA results in host gene knock-down, which is partially rescued upon host *Dicer* knock-down** (**A**) qPCR of mRNA extracted from day 3 of chimera-RNAi feeding, showing change in gene expression of *EF1-α*, *HSP90* and *tub-β* gene in *P. bursaria* in response to chimera dsRNA exposure (yellow/gold), compared to *Dicer* dsRNA mixed controls (grey). Change of relative expression (ddCT) was calculated for treated (chimera dsRNA or chimera/Dicer dsRNA) vs untreated (scramble dsRNA) control cultures, and normalised against the standardised change in expression of an Actin housekeeping gene. Multiple vector delivery was conducted at a 50:50 ratio during feeding. Data are represented as mean ± SEM of three biological replicates. This figure is an extension of **Figure 3E**, demonstrating how reduction in host transcript expression in response to endosymbiont chimera dsRNA exposure is partially rescued by host Dicer knock-down.

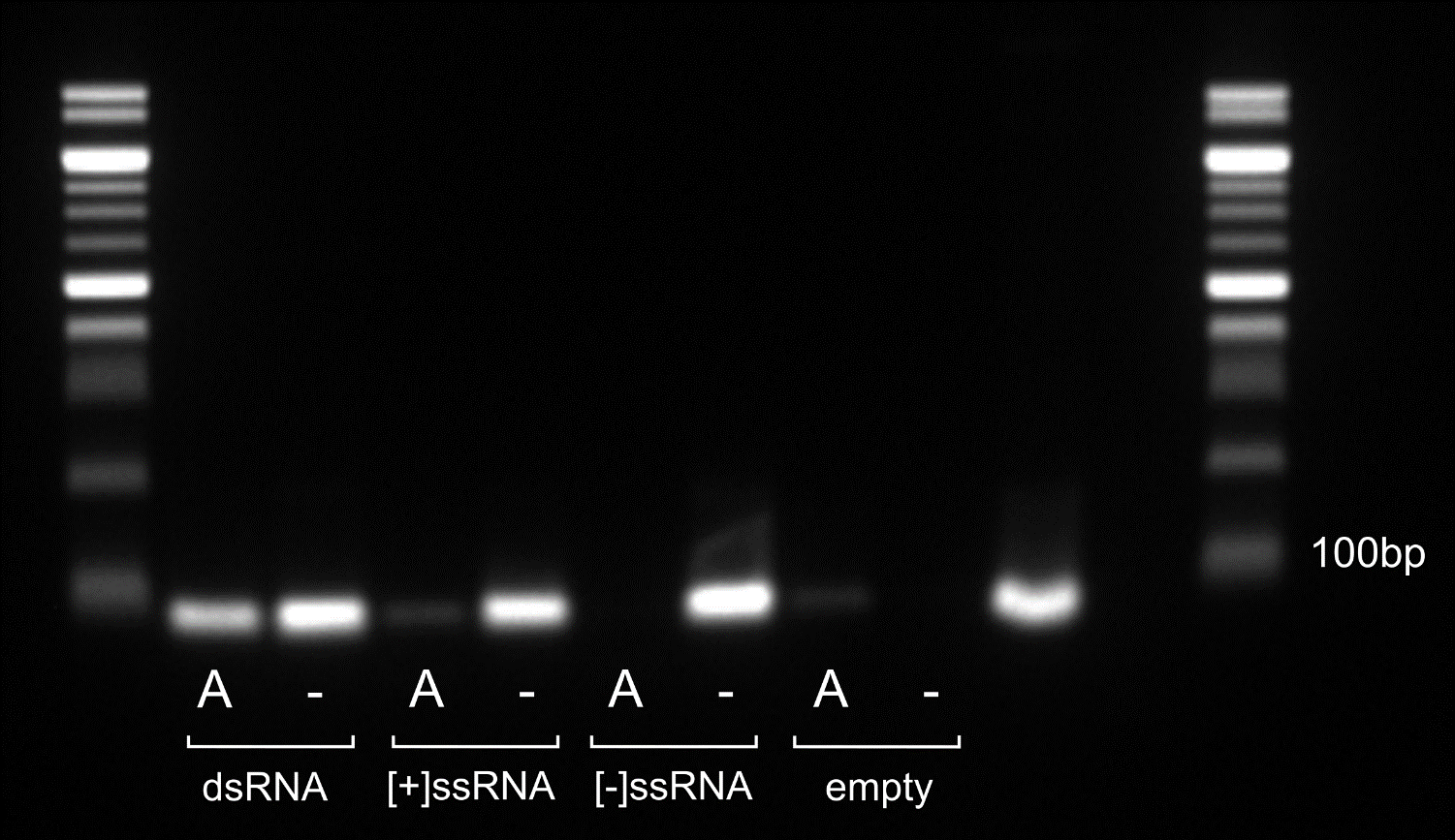

**Figure S13. Validation of single-stranded RNAi vector construction, confirming the production of ssRNA.** Gel image of an 85-bp reverse transcriptase PCR product produced from an L4440 construct expressing dsRNA, sense [+]ssRNA, or antisense [-]ssRNA, compared to an empty vector control. Treatment with RNAse A (A) was used to degrade ssRNA, leaving only dsRNA. Untreated samples (-) would still contain ssRNA. Wells 9 and 10 indicate positive and negative template controls, respectively. This data confirms the production of ssRNA in both sense and antisense orientation by the single-stranded vector constructs used in **Figure 4**.

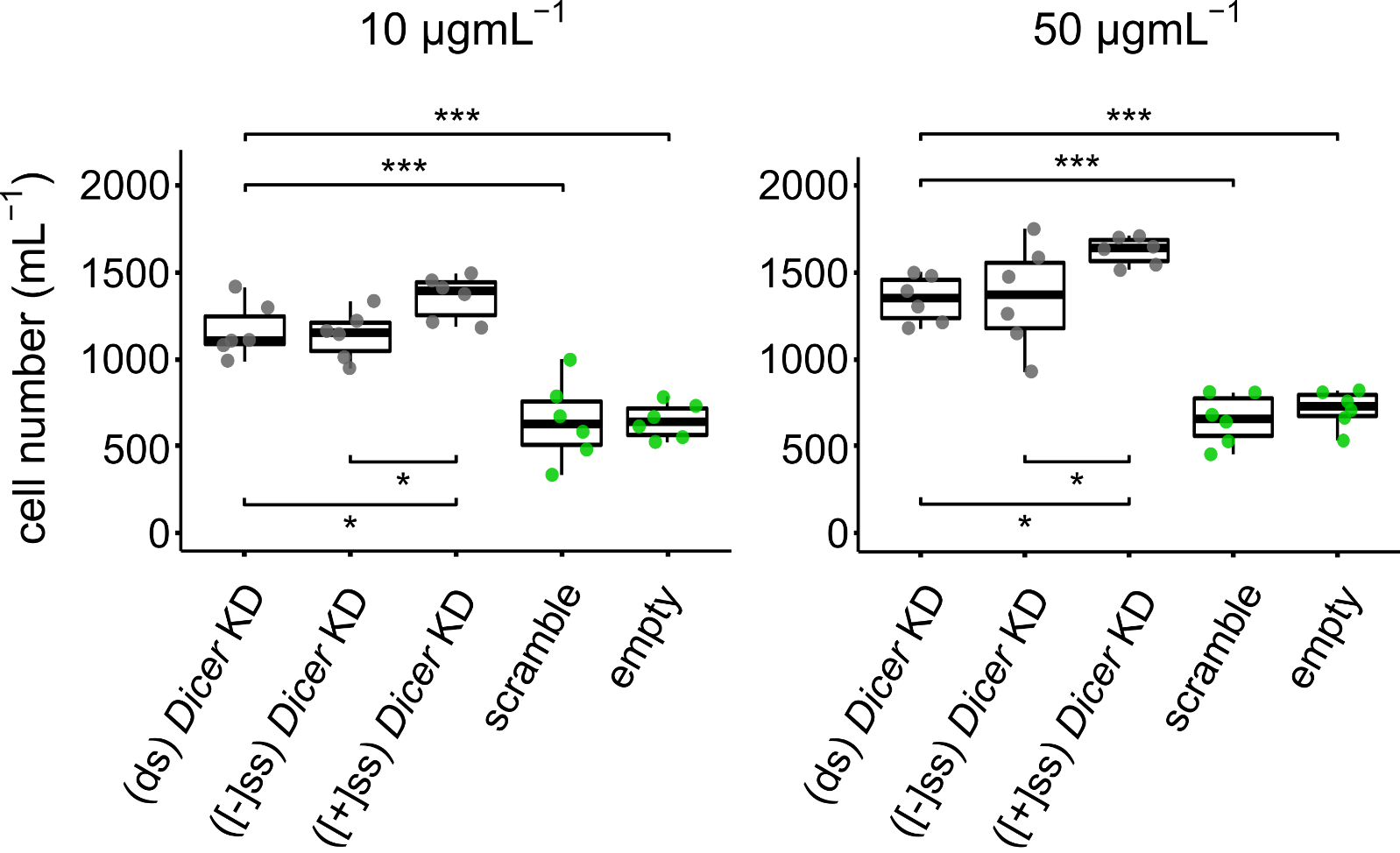

**Figure S14. Knockdown of host *Dicer* function using *Dicer* ssRNA (in the sense orientation) provides greater rescue efficacy than knockdown using *Dicer* dsRNA, during cycloheximide treatment, and demonstrates that sense ssRNA is the most potent substrate for RNAi induction.** *P. bursaria* cell number after 8 days of treatment with cycloheximide at 10 µgmL^-1^ or 50 µgmL^-1^. *P. bursaria* cultures were simultaneously fed for 12 days (starting four days prior to drug treatment) with *E. coli* expressing; *Dicer* dsRNA, *Dicer* antisense [-]ssRNA, *Dicer* sense [+]ssRNA, non-hit ‘scramble’ dsRNA, or an empty vector control. Note the relative effect of *Dicer* [+]ssRNA exposure compared to *Dicer* dsRNA exposure, and compared to scramble and empty vector controls. Boxplot data are represented as max, upper quartile (Q3), mean, lower quartile (Q1) and min values of six biological replicates. Significance calculated as *p ≤ 0.05, ***p ≤ 0.001 using a generalized linear model with quasi-Poisson distribution. This figure is an extension of **Figure 1E** showing that RNAi function, mediated by *Dicer*, results in a cost to growth during endosymbiont elimination. These data also relate to **Figure 4**, which demonstrates the increased potency of [+]ssRNA delivery to induce an RNAi-mediated effect.

**Table S1. Full list of mRNA-based constructs,** used for feeding-based RNAi experiments in this study. See attached Excel spreadsheet.

**Table S2. eDicer RNA-RNA interaction predictions**, used for the data analysis in **Figure S8**. See attached Excel spreadsheet.

**Table S3. Endosymbiont mRNA-mRNA interaction fragments,** used to create the synthetic endosymbiont derived chimera in **Figure 3.** See attached Excel spreadsheet.

**Table S4. SSU rRNA derived putative rRNA-rRNA ‘collision’ fragments,** used in **Figure S9**. See attached Excel spreadsheet.

**Table S5. Full list of primers and conditions,** used for qPCR experiments in this study. See attached Excel spreadsheet.

**SUPPLEMENTARY RESULTS AND DISCUSSION**

**Discussion S1**

***Considerations for synthetic endosymbiont derived RNA exposure to simulate naturally occurring RNA interactions.***

It is important to consider that delivery of a synthetic endosymbiont-derived RNA chimera in this manner represents only an approximation of the putative 23-nt mRNA-mRNA interactions that may be occurring in the *P. bursaria*-*Chlorella spp.* system. Delivery of synthetic endosymbiont derived RNA via an *E. coli* vector, as outlined in **Figure 3** and **Figure 4**, may over-inflate the nature of any identified mRNA-mRNA interactions relative to those naturally derived from the endosymbiont. Upon delivery of the 450-nt synthetic endosymbiont derived chimera, cleavage via Dicer would result in the generation of 428 possible unique 23-nt outcomes (450 – 23 + 1). As such, the manner in which these ten endosymbiont derived mRNA interaction fragments are arranged within the chimera would mean that only 2.3% (10/428) of all generated siRNAs could result in an interaction with host mRNA. This compares to the 0.121% predicted interaction rate between endosymbiont-and-host transcripts (**Figure S8A**).

This predicted chimera interaction rate of 2.3% would be further diluted by RNA derived from the *E. coli* feeding vector. We also note that knock-down of only three of the targeted transcripts (*EF-1α, HSP90 and tub-β*) was capable of inducing a cost to host growth when the wider corresponding host transcript was targeted directly (**Figure 3D;** see also **Figure S11**). Importantly, we cannot rule out the potential for synergistic effects of numerous simultaneous knock-downs, which may amplify a cumulative negative effect on host growth dynamics. It can also be anticipated that these synergistic effects would be further amplified by exposure to a wider eukaryotic endosymbiont mRNA population from multiple endosymbiotic algae. Furthermore, a 1:3 dilution of chimera dsRNA delivery alongside a ‘nonsense’ encoding vector was also able to induce a cost to host growth (**Figure S9**). These results indicate that a relatively small quantity of mRNA-mRNA interactions may be sufficient to induce a cost to host growth in *P. bursaria*.

***Putative mechanistic explanation for a sense oriented ssRNA knock-down bias***

The basis of a putative RNA orientation bias during induction of RNAi has not yet been investigated in *Paramecium*. In fact, a feeding-based approach to RNAi induction through delivery of ssRNA has been previously deemed ineffective in *P. tetraurelia*^2–4^. Functional asymmetry in siRNA is a phenomenon in which sequence composition of each strand of the siRNA duplex, particularly stability of the 5’ ends, has been shown to determine the degree to which each strand is loaded onto the RISC to direct complementary mRNA cleavage^5–7^. This functional asymmetry forms the basis of miRNA biogenesis in other systems, in which the miRNA of a transient double-stranded precursor is preferentially selected to enter the RNAi-induced silencing complex, resulting in the accumulation of miRNA *in vivo* as ssRNA^7^. *Paramecium* does not appear to produce miRNA, however preliminary evidence suggests that these may be functionally encapsulated by an un-derived siRNA-based RNAi pathway capable of functioning in endogenous transcriptome regulation^8,9^.

Due to the potential for functional asymmetry, it should be considered therefore that efficacy of ssRNA may not necessarily be derived from orientation of delivery, but rather resultant stability of the RdRP- and Dicer- generated siRNA duplex. In the above examples, stability preference was demonstrated through delivery of the siRNA duplex rather than dsRNA, which would be randomly cleaved into many siRNA duplexes of variable stability. If this mechanism were responsible for the orientation bias observed in **Figure 4** & **Figure S14**, then a potential explanation would be that this bias is due to the outcome of RdRP-mediated generation of duplex siRNA from a single-stranded precursor, resulting in preferential loading of (for example) the newly generated strand onto the RNAi-induced silencing complex^7,10^. Under this hypothesis, both strands may go on to induce silencing, however delivery of [+]ssRNA would potentially induce silencing at a greater efficacy than dsRNA (a 50:50 dilution of both sense and antisense ssRNA) or [-]ssRNA (which would result in generation and preferential loading of the non-complementary strand, although over time this could still generate the same response given sufficient iterations). Nonetheless, these explanations remain speculative and require a far greater understanding of the pathways and factors that facilitate ssRNA-induced RNAi in *P. bursaria* to be elucidated in full.

**SUPPLEMENTARY METHODS**

***Full overview of the eDicer comparative analysis process***

Using a dataset consisting of transcripts binned as either ‘endosymbiont’ or ‘host’ (derived from a published *P. bursaria* transcriptome^11^), we sought to identify all of the possible endosymbiont-host mRNA-mRNA interactions that could occur in the *P. bursaria*-algal system. These represented endosymbiont derived sequences which, if processed by the host Dicer endonuclease into ~23-nt siRNA^12^, could retain sufficient sequence identity to putatively knock-down expression of host genes via the RNAi mediated mechanism. Due to the potential bidirectional transcription of ssRNA into dsRNA by host RNA-dependent RNA polymerase (RdRP) prior to Dicer processing, the eDicer and mapping processes considered both sense and antisense complementation between each RNA population. Using this approach, we identified 35,703 distinct 23-nt putative mRNA-mRNA interactions between the *P. bursaria* host and algal endosymbiont transcript datasets, representing 0.121% of the total inventory of distinct host 23-nt k-mers identified (**Figure S8A** & **Table S2**).

Two further datasets were compared to the host, each containing genes predicted from bacterial genomic data. These were assessed in order to identify the number of putative mRNA-mRNA interactions derived from bacterial food acquired during culturing (*Klebsiella pneumoniae* [NCBI acc.: NC_016845.1]^13^) or vector-based RNAi feeding (*Escherichia coli* [NCBI acc.: NZ_CP012868.1]^14^). In this manner, we identified that the proportion of distinct 23-nt putative mRNA-mRNA interactions between endosymbiont-and-host transcripts was proportionally 120-fold greater than between these bacterial food-and-host transcripts (**Figure S8A** & **Table S2**).

We note that a greater number of putative mRNA-mRNA interactions between endosymbiont-and-host are possibly produced from a relatively small number of extended regions of sequence identity shared between conserved eukaryotic transcripts. Notably, these regions of shared sequence identity would be absent in the prokaryote-eukaryote transcript comparison between bacterial food-and-host, resulting in the large-scale increase in the number of putative mRNA-mRNA interactions predicted between *P. bursaria* and endosymbiont (i.e. a 120-fold increase). Parallel comparisons of the 21-22-23 nt interactions generated by eDicer showed a relatively stable proportion of putative mRNA-mRNA interactions between endosymbiont-and-host (eukaryote-eukaryote) across these sRNA sizes (i.e. 0.13/0.125/0.121% for the respective 21-22-23 nt fragment comparisons). In contrast, the relative proportion of putative mRNA-mRNA interactions between bacterial food-and-host (prokaryote-eukaryote) showed a greater degree of variation (i.e. 0.016/0.004/0.001% for *E. coli* & 0.011/0.003/0.001% for *K. pneumoniae*, for each respective 21-22-23 nt fragment comparison), consistent with the idea that a short extension in shared sequence identity between eukaryotic transcripts of different sources represents an important factor for generating a higher relative rate of RNA-RNA interaction. Nonetheless, these data indicate that the number of putative mRNA-mRNA interactions predicted between endosymbiotic algal and host mRNA populations is greater than those predicted between bacterial food and host mRNA populations.

All datasets were cross referenced to a yeast ‘lethal gene’ database^15^ which contained genes known to be conditionally essential in *Saccharomyces cerevisiae* (1 in 5.5 of the total yeast genes investigated)*.* These curated ‘lethal’ datasets were then compared using eDicer in order to identify the proportion of putative mRNA-mRNA interactions that could result in knock-down of homologues of these conditionally essential yeast genes in *P. bursaria,* and thus could be considered to be ‘putatively lethal’. Using this approach, we identified 4,776 distinct 23-nt putative ‘lethal’ mRNA-mRNA interactions between endosymbiont-and-host, representing 0.168% of the total inventory of distinct ‘lethal’ host 23-nt k-mers (**Figure S8A** & **Table S2**). Once again, this value was considerably greater than the number of putative ‘lethal’ mRNA-mRNA interactions predicted between the bacterial food-and-host transcript datasets.

Importantly, these data demonstrate that a considerable proportion (1 in 7.5) of the total distinct 23-nt putative mRNA-mRNA interactions predicted between endosymbiont-and-host involved putatively ‘lethal’ host transcripts. For reference, only 1 in 10.4 distinct host 23-nt k-mers overall were classified as homologues of these essential yeast genes and thus were considered to be ‘lethal’. Within the distinct 23-nt fragment dataset, this suggests a ratio of total:‘lethal’ putative mRNA-mRNA interactions of 1:1.4 (% of ‘lethal’ mRNA-mRNA interactions / % of total mRNA-mRNA interactions). This contrasts with the ratio of total:‘lethal’ putative mRNA-mRNA interactions predicted between bacterial food-and-host of 1:0.16 (*E. coli*) and 1:0.17 (*K. pneumoniae*). These comparisons therefore demonstrate that not only is there a greater proportion of putative mRNA-mRNA interactions between endosymbiont-and-host within the ‘lethal’ dataset (compared to the total transcript dataset), but also that these putative mRNA-mRNA interactions between endosymbiont-and-host (eukaryote-eukaryote) are more likely to be enriched for transcripts of putatively ‘lethal’ genes, compared to between bacterial food-and-host (prokaryote-eukaryote).

Concurrently, a further dataset was curated consisting of full-length ribosomal RNA (rRNA) clusters for *Microactinium conductrix* (a *P. bursaria* endosymbiont [NCBI acc.: ASM224581v2]^16^), *K. pneumoniae* (food [NCBI acc.: NC_016845.1]^13^), *E. coli* (vector-based RNAi food [NCBI acc.: NZ_CP012868.1]^14^) and two strains of host *P. bursaria* – Yad1g1N (modified from published transcriptome data^11^) and 186b (identified from our sequence data). These were once again compared using eDicer in order to identify all of the possible rRNA-rRNA interactions that could occur in the *P. bursaria*-algal system. We observed an approximately 200-to-230-fold increase (186b and Yad1g1N respectively) in the number of distinct 23-nt putative rRNA-rRNA interactions between endosymbiont-and-host compared to bacterial food-and-host (**Figure S8B** & **Table S2**). Collectively, these *in silico* predictions demonstrate that there is far greater potential for the occurrence of both mRNA-mRNA and rRNA-rRNA interactions between the algal endosymbiont and host (eukaryote-eukaryote) RNA populations, than there is between consumed bacterial food and host (prokaryote-eukaryote) RNA populations. Importantly, for putative mRNA-mRNA interactions predicted between two eukaryotic RNA populations (such as between *P. bursaria* and endosymbiotic algae), these were also more likely to be enriched for transcripts of putatively essential (‘lethal’) host genes.
